# Laboratory strains of *Escherichia coli* K-12: not such perfect role models after all

**DOI:** 10.1101/2022.06.29.497745

**Authors:** Douglas F. Browning, Jon L. Hobman, Stephen J.W. Busby

## Abstract

*Escherichia coli* K-12 was originally isolated 100 years ago and since then, it has become an invaluable model organism and a cornerstone of molecular biology research. However, despite its apparent pedigree, since its initial isolation, *E. coli* K-12 has been repeatedly cultured, passaged, and mutagenized, resulting in an organism that carries extensive genetic changes. To understand more about the evolution of this important model organism, we have sequenced the genomes of two ancestral K-12 strains, WG1 and EMG2, considered to be the progenitors of many key laboratory strains. Our analysis confirms that these strains still carry genetic elements such as bacteriophage lambda (λ) and the F plasmid, but also indicates that they have undergone extensive lab-based evolution. Thus, scrutinizing the genomes of ancestral *E. coli* K-12 strains, leads us to question whether *E. coli* K-12 is a sufficiently robust model organism for 21st century microbiology.

**DATA SUMMARY:** All supporting data are provided within the article or through supplementary data files. Supplementary Figs. S1 to S14 and Supplementary File S1 are available with the online version of this article. All genome sequence data has been deposited in NCBI GenBank under Bioproject ID PRJNA848777. The assembled and annotated genomes of WG1 and EMG2 have been deposited with the accession numbers, CP099590 and CP099591 (WG1) and CP099588 and CP099589 (EMG2).

**Impact Statement:** Since its isolation in 1922, *Escherichia coli* K-12, has become arguably the premier model organism for contemporary science. The adoption of *E. coli* K-12 by many microbiologists across the globe, means that it has a complex pedigree, and, although many *E. coli* K-12 strains have been sequenced, little is known about the early versions of K-12, which still carry the F plasmid and bacteriophage λ. To understand more about the lab-based evolution that has shaped this important model organism, we have sequenced two ancestral K-12 strains, WG1 and EMG2, that are considered to be the progenitors of many of the laboratory strains used today.

## INTRODUCTION

*Escherichia coli* K-12 was originally isolated in 1922 from a convalescent diphtheria patient and, later in the 1940s, adopted by Charles Clifton and Edward Tatum as a model organism (1-3). Since then, *E. coli* K-12 has become the “workhorse” of molecular biology, becoming arguably the premier model organism in science today. MG1655 was the first *E. coli* K-12 strain to have its genome sequence published, followed by W3110, resulting in an explosion of genomic research and comparative genomics (4, 5). However, despite its prestige, *E. coli* K-12 was stored on agar plates, stabs or slopes before cryopreservation became established, and has been repeatedly subcultured and mutagenized (Fig. 1), resulting in an organism which carries extensive genetic changes and has lost the ability to produce many surface-associated structures (3). For example, *E. coli* K-12 lab strains are unable to synthesize O antigen on their lipopolysaccharide and no longer carry the F plasmid or bacteriophage λ (3, 6-9). One major strength of using *E. coli* K-12 strains for cloning and heterologous gene expression is that K-12 strains cannot establish in the human gut (10, 11), and, thus, even so-called “wild type” *E. coli* K-12 strains, like MG1655 and W3110, are very different from commensal or environmental isolates (3, 4, 12, 13). To understand more about the evolution of this important model organism, we have sequenced the genomes of two *E. coli* K-12 strains, WG1 and EMG2, the proposed ancestors of key laboratory strains (Fig. 1) (1, 2). Our analysis confirms that these strains carry genetic elements such as phage λ and the F plasmid, but indicates that they have also undergone extensive mutational alternation during their evolution in laboratories.

**Fig. 1.**
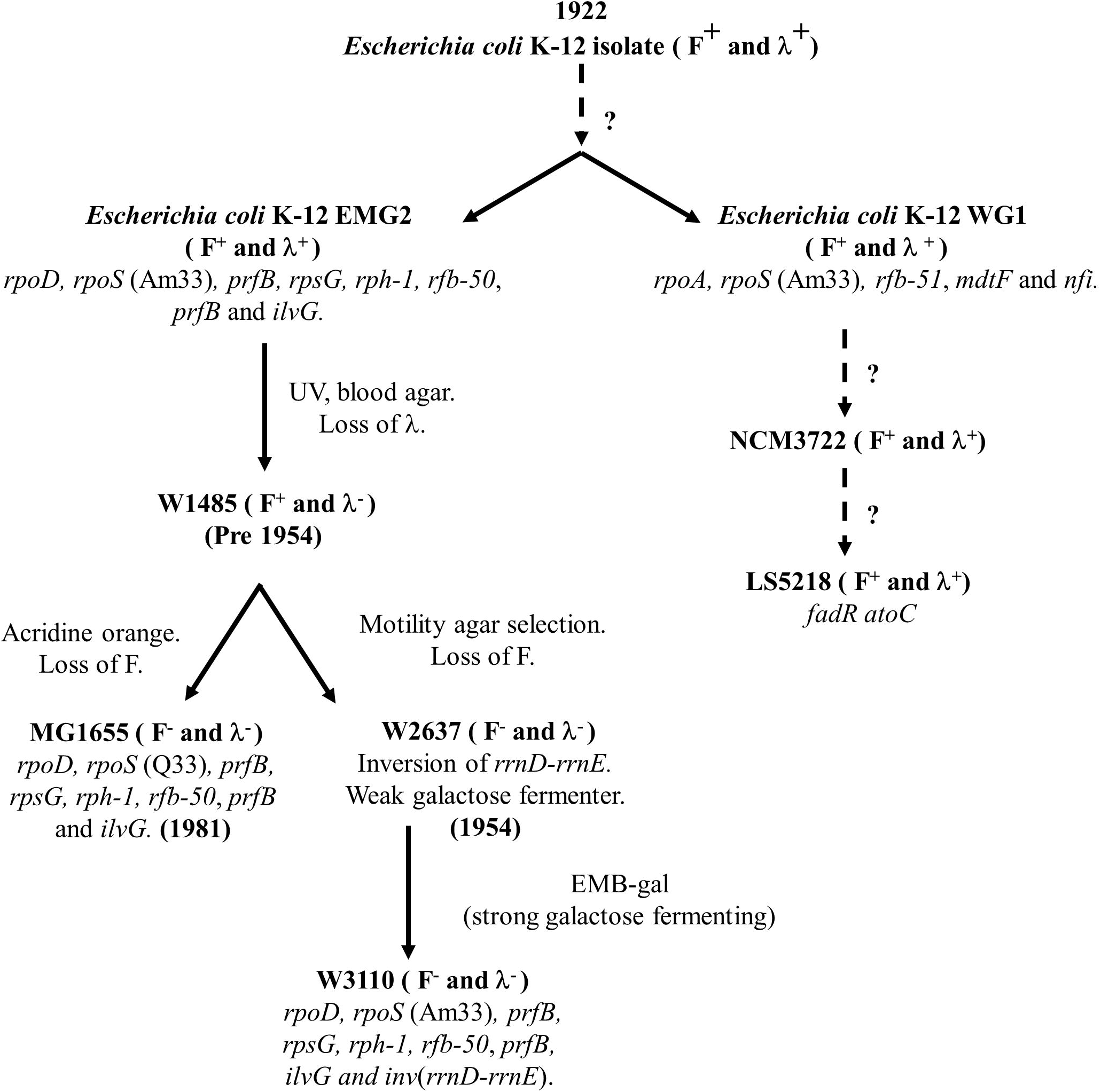
The pedigree of *Escherichia coli* K-12 strains. The figure details the pathway of *E. coli* K-12 evolution from its isolation in 1922 to the generation of MG1655 and W3100 strains (1, 3, 4, 12). Blood agar indicates selection on blood agar plates; UV, irradiation with ultraviolet light; EMB-gal, selection for utilization of galactose on eosin methylene blue indicator plates. Dotted lines represent uncertain evolutionary lineage events.

## METHODS

### Bacterial strains and whole genome sequencing

*E. coli* K-12 strains WG1 and EMG2 were obtained for the Coli Genetic Stock Centre (CGSC), strain numbers CGSC#5073 and CGSC#4401, respectively (1, 2). Each strain was sequenced using the enhanced sequencing option from MicrobesNG (https://microbesng.com/), which uses a combination of Illumina and Oxford Nanopore Technologies (ONT). Cell cultures were grown in LB medium and the cell pellet was isolated by centrifugation and resuspended in the cryo-preservative in a Microbank™ tube (Pro-Lab Diagnostics UK, United Kingdom). Approximately 2×10^9^ cells were used for high molecular weight DNA extraction using Nanobind CCB Big DNA Kit (Circulomics, Maryland, USA). DNA was quantified with the Qubit dsDNA HS assay in a Qubit 3.0 (Invitrogen). Long read genomic DNA libraries were prepared with the Oxford Nanopore SQK-LSK109 kit with Native Barcoding EXP-NBD104/114 (ONT, UK), using 400-500 ng of high molecular weight DNA. Twelve to twenty-four barcoded samples were pooled in a single sequencing library and loaded on a FLO-MIN106 (R.9.4 or R.9.4.1) flow cell in a GridION (ONT, UK). Illumina reads were adapter trimmed using Trimmomatic 0.30 with a sliding window quality cutoff of Q15 (14). Unicycler v0.4.0 was used for genome assembly (15) and Prokka 1.11 to annotate contigs (16). Sequence data has been deposited at DDBJ/ENA/GenBank with the accession numbers CP099590 and CP099591 for WG1 and CP099588 and CP099589 for EMG2.

### Bioinformatic analysis of genome sequences

For single nucleotide variant (SNV) calling, reads from EMG2 were aligned to the WG1 reference genome using BWA-Mem and processed using SAMtools 1.2. Variants were called using VarScan with two thresholds, sensitive and specific, where the variant allele frequency is greater than 90% and 10% respectively. The effects of variants were predicted and annotated using SnpEff. Draft genomes were visualized using Artemis (17), and comparisons between *E. coli* K-12 genomes were made using the Basic Local Alignment Search Tool (BLAST) at NCBI (https://blast.ncbi.nlm.nih.gov/Blast.cgi), the Artemis Comparison Tool (ACT) (18) and the Proksee Server (https://proksee.ca/) (19). Genome representations were drawn using the Proksee Server (19) and ACT (18). Plasmid replicons were detected in draft genomes with PlasmidFinder 2.1 (5), using the software at the Center for Genomic Epidemiology (CGE) (http://www.genomicepidemiology.org/). Insertion sequences were located using ISfinder (https://www-is.biotoul.fr/blast/resultat.php) (20).

## RESULTS

### Comparison of the WG1 and EMG2 genomes

Whole genome sequencing of WG1 and EMG2 resulted in draft genome sequences, each comprising of 2 contigs; the larger contig, Contig 1, is the chromosomal sequence and the smaller, Contig 2, is the F plasmid (Figs. 2 and 3, Supplementary Fig. S1: Tables 1 and 2). Since both strains carry bacteriophage λ and the F plasmid, their genomes are slightly bigger than other sequenced *E. coli* K-12 strains, such as MG1655 and W3110 (Table 1) (4, 12). Comparison of the genomes of both WG1 and EMG2 with those of MG1655 and W3110 indicated that, unlike W3110, no major chromosomal rearrangements had occurred in these strains (Supplementary Fig. S2) (19, 21). However, we identified a number of obvious regions of difference (Fig. 2 and Supplementary Fig. S1 and S3). For example, both EMG2 and W3110 have lost the cryptic prophage CPZ-55, and EMG2 has lost the *gatYZABDR* locus, which is involved in galactitol metabolism (22) (Fig. 2 and Supplementary Fig. S4). Interestingly, the *gatYZABDR* genes appear to have been a hotspot for insertion sequence element disruption in both MG1655 and W3110, which affects expression of this region (Supplementary Fig. S5) (22). Similarly, the region upstream of *flhDC* locus, which controls flagella production, also seems to have been targeted by different transposable elements (Fig. 2 and Supplementary Fig. S6) (23, 24). Note that strains that have been stored in agar stabs for many years accumulate deleterious mutations due to wholesale transposition of insertion sequences (25-27). As insertion of elements into this region influences motility, it is likely that the sequence heterogeneity found in this region produces a spectrum of effects (23, 24). For WG1, we detected the loss of cryptic prophage CP4-6 and a large deletion of the lipopolysaccharide O-antigen biosynthetic cluster, previously termed *rfb-*51 (Supplementary Figs. S1, S3 and S4) (8). Note that EMG2, MG1655 and W3110 carry the alternative *rfb-*50 mutation (an IS*5* disruption of the rhamnose transferase gene *wbbL*), which appears to be common to most *E. coli* K-12 strains (28), and so do not produce O-antigen either (Supplementary Figs. S4c) (8, 9). Loss of O-antigen production seems likely to be an adaptation to laboratory life, with both the first *E. coli* strain NCTC 86 (isolated in 1885), and commonly used B strains (*e*.*g*., BL21(DE3)), all being rough in nature (13, 29, 30). In addition to these differences, WG1 also carries additional genes, encoding an LPS export ABC transporter permease (*lptG*), an acyl-carrier protein (*acpP*) and a NAD-dependent epimerase/dehydratase (*oleD*), which are flanked by IS*5* elements (Fig. 2 and Supplementary Fig. S7).

**Fig. 2.**
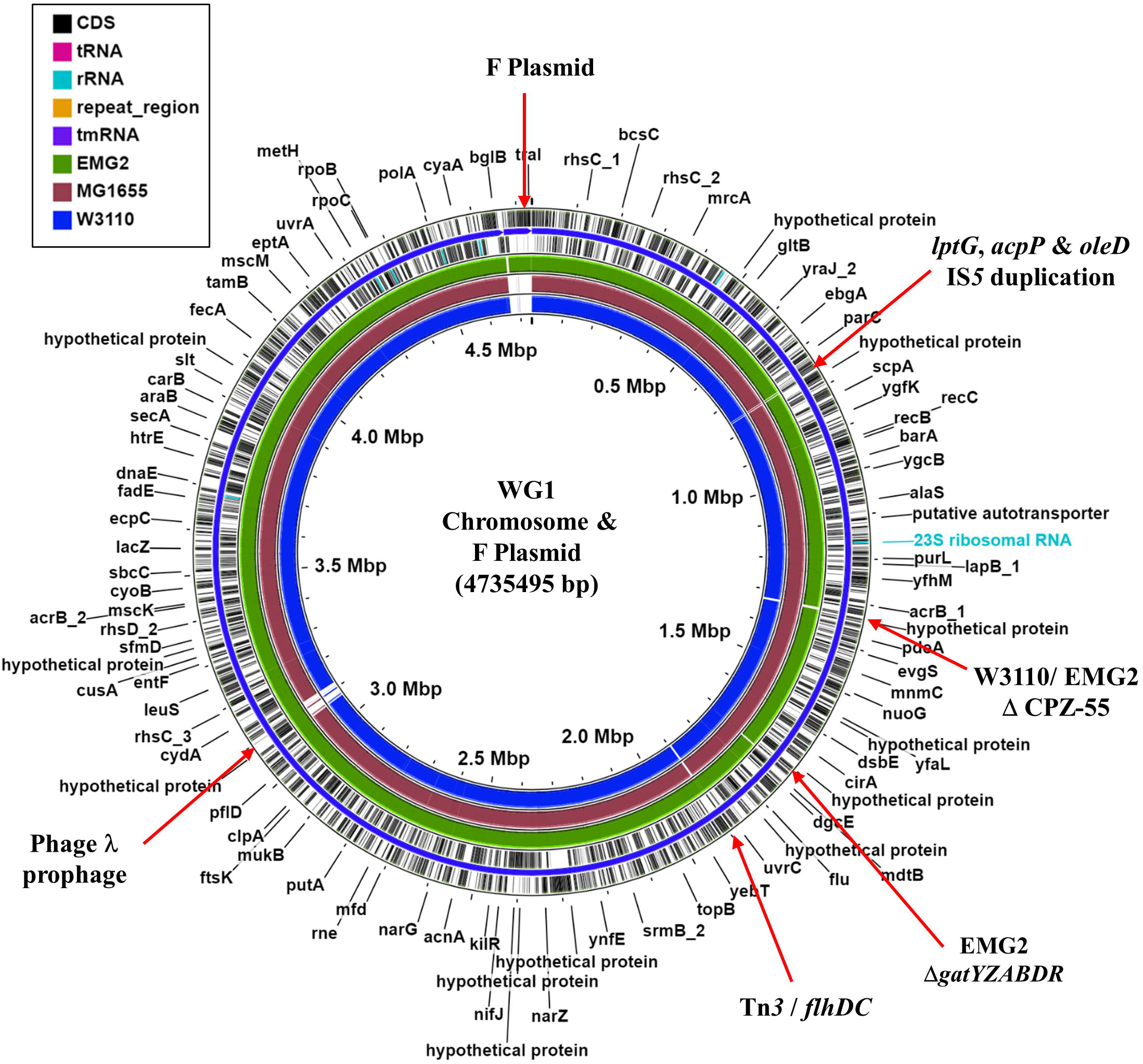
Genome comparison of different *E. coli* K-12 strains. The figure shows the comparison of the WG1 chromosome (contig 1) and F plasmid (contig 2) with the genomes of EMG2, MG1655 (NC_000913.3) and W3110 (NC_007779.1), using the Proksee Server (19). The outer two rings display the genes and features of the WG1 genome, with selected genes and differences labelled. The green, brown and blue rings illustrate the BLAST results when the genome sequences of *E. coli* K-12 strains EMG2, MG1655 (NC_000913.3) and W3110 (NC_007779.1), respectively, are compared to that of WG1.

**Fig. 3.**
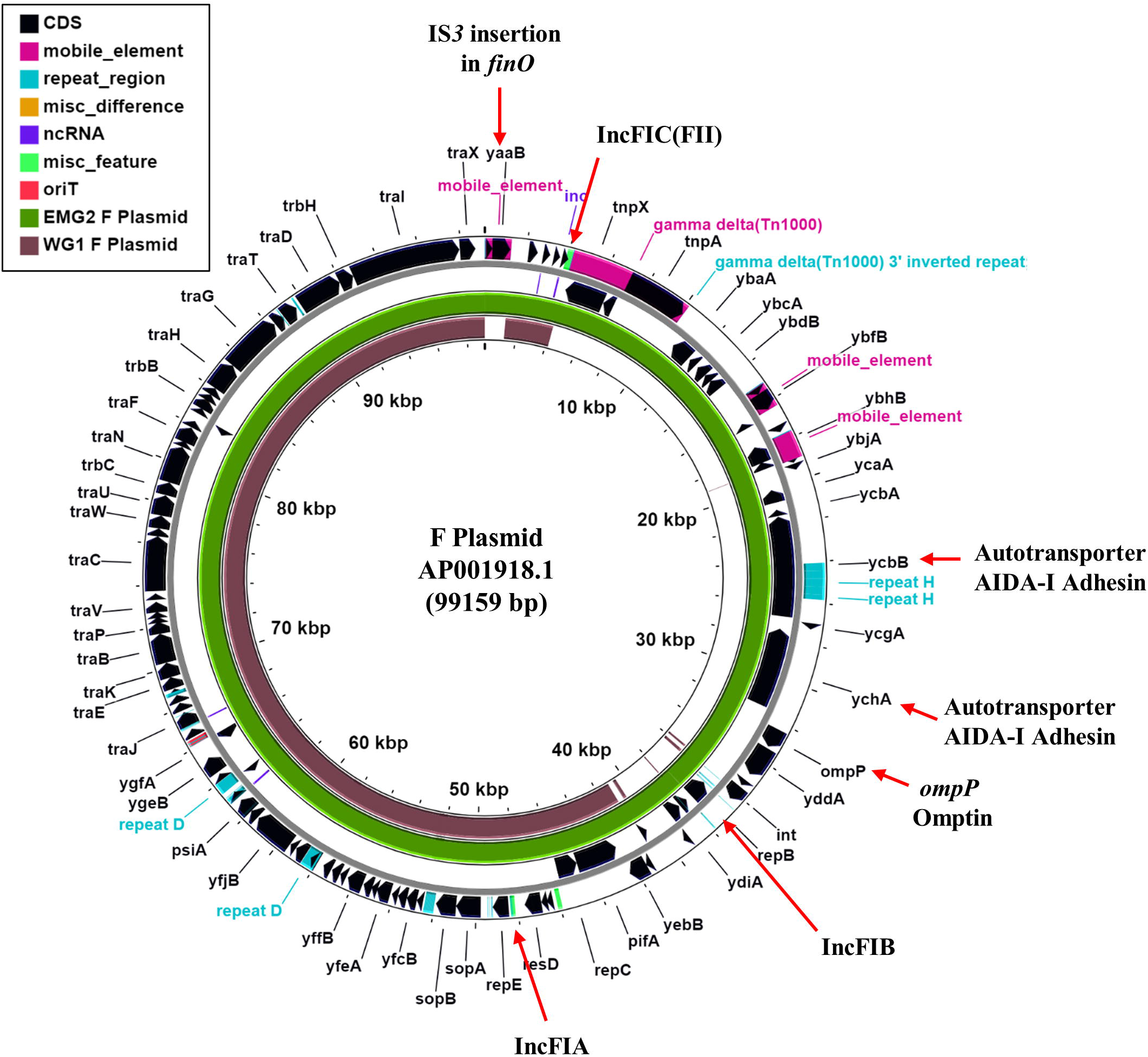
Comparison of the F plasmids from different *E. coli* K-12 strains. The figure shows the comparison of the F plasmid (AP001918.1) with that from EMG2 and WG1 using Proksee (19). The outer two rings display the genes and features of the F plasmid, with selected genes labelled. The green and brown rings illustrate the BLAST results when the F plasmid sequences from EMG2 and WG1, respectively, are compared to the original F plasmid sequence.

**Table 1.** Comparison of the genomes of different *E. coli* K-12 laboratory strains.

|  | **WG1** | **EMG2** | **MG1655** | **W3110** | **NCM3722** | **LS5218** |
| --- | --- | --- | --- | --- | --- | --- |
| Accession N° | CP099590<br>CP099591 | CP099588<br>CP099589 | NC_000913.3 | NC_007779.1 | CP011495.1<br>CP011496.1 | MVJG000000<br>00.1 |
| Genome size <sup>a</sup> | 4,735,495 | 4,774,480 | 4,641,652 | 4,646,332 | 4,745,591 | 4,699,198 |
| Plasmid | F plasmid | F plasmid | None | None | F plasmid | F plasmid |
| Total N° CDSs <sup>b</sup> | 4431 | 4457 | 4285 | 4213 | 4539 | 4368 |
| GC content | 50.75 % | 50.73 % | 50.79 % | 50.8 % | 50.76 % | 50.72 % |

<sup>a</sup> Genome size (bp) includes the F plasmid for WG1, EMG2, NCM3722 and LS5218.
<sup>b</sup> Number of coding sequences (CDSs) is as predicted by each genome annotation.

**Table 2.** Comparison of the F plasmid from different *E. coli* K-12 laboratory strains.

|  | **WG1** | **EMG2** | **F Plasmid** | **NCM3722** | **LS5218** |
| --- | --- | --- | --- | --- | --- |
| Accession | CP099591 | CP099589 | AP001918.1 | CP011496.1 | CM007715.1 |
| F Plasmid size | 67,408 bp | 99,158 bp | 99,159 bp | 67,545 bp | 67,502 bp |
| Total N° CDSs <sup>a</sup> | 73 | 98 | 105 | 79 | 83 |
| GC content | 51.66 % | 48.17 % | 48.17 % | 51.67 % | 51.67 % |
| Plasmid Replicons <sup>b</sup> | IncFIA,<br>IncFIC(FII) | IncFIA,<br>IncFIB,<br>IncFIC(FII) | IncFIA,<br>IncFIB,<br>IncFIC(FII) | IncFIA,<br>IncFIC(FII) | IncFIA,<br>IncFIC(FII) |

<sup>a</sup> Number of CDSs is as specified by genome annotation.
<sup>b</sup> Plasmid replicons were detected using PlasmidFinder 2.1 using software at the CGE (5).

### The F Plasmid

Comparison with MG1655 confirmed that both WG1 and EMG2 both carry the F plasmid, however, the two versions of F differ markedly in size, with that from EMG2 (99158 bp) similar in size to the previously sequence F plasmid (AP001918.1: 99159 bp), whilst F from WG2 is considerable smaller (67408 bp) (Table 2: Fig. 3 and Supplementary Fig. S8). This can be attributed to the loss of a large section of F in WG1, carrying the AIDA-I like autotransporter adhesin genes *ycbB* and *ychA*, the *ompP* omptin and the IncFIB replicon (Table 2: Fig. 3 and Supplementary Figs. S9) (7, 31). Surprisingly, F from WG1 carries additional DNA that is not found on F, which includes an IncFII RepA protein (Supplementary Figs. S8 and S9). As in the previously sequenced F plasmid (AP001918.1), EMG2 F carries an IS*3* insertion in the *finO* gene, which leads to constitutive F transfer (7, 32, 33). However, this insertion sequence is absent from the WG1 F (Fig. 3; Supplementary Figs. S8 and S9), suggesting that conjugative transfer is regulated in this plasmid and that the insertion of IS*3* must have occurred in the immediate ancestor of EMG2. Thus, it is clear that F plasmids from both EMG2 and F have undergone significant lab-based evolution, resulting in two very different plasmids.

### Bacteriophage λ

Comparison with MG1655 indicated that, as expected, both WG1 and EMG2 carry the bacteriophage λ prophage integrated between the *bioA* and *ybhC* genes (Fig. 2 and Supplementary Figs. S1 and S10). However, comparison with the previous sequenced λ genome (NC_001416) identified some differences in λ from WG1 and EMG2, in particular with the genes encoded tail fibres J, Stf and Tfa (Supplementary Fig. S11). Of note is *stf* (side tail fibre), which, in λ (NC_001416), carries a frame shift disrupting the gene into two ORFs (*orf-401* and *orf-314*) (34, 35). Bacteriophage λ carrying this lesion (λ PaPa) forms larger λ plaques (6, 35). Thus, as *stf* remains intact in WG1 and EMG2, it is likely that both strains would produce a small plaque phenotype (6, 35).

### Similarities and differences between WG1 and EMG2

Single nucleotide variant calling showed that *E. coli* K-12 strains WG1 and EMG2 also differ in a number of key genes involved in important cellular functions (Supplementary File S1). For example, in EMG2, the gene encoding the major sigma factor σ^70^ (*rpoD*), carries a substitution, which results in Try at position 571 (Supplementary Fig. S12a). This is also found in MG1655 and W3110, whilst most *E. coli* strains carry His at this position. Substitutions at σ^70^ residue 571 have been shown to affect transcription at the *lac, araBAD, merT, merR* and the P22 phage *ant* promoters, as well as interfering with σ^70^ binding to core RNA polymerase and its ability to compete with alternative sigma factors (36-40). Conversely, in WG1 the gene encoding the α subunit of RNA polymerase, carries a mutation which results in a Gly to Arg substitution at position 311 (Supplementary Fig. S12b). This alteration affects expression from both the *merT* and *merR* promoters and the anaerobically activated *pepT* promoter in *Salmonella enterica* serovar Typhimurium (38, 41) (note that α in *E. coli* and *S. enterica* serovar Typhimurium are identical). As for many K-12 strains, both WG1 and EMG2 carry a truncation in *rpoS*, which encodes the stress and stationary phase sigma factor σ^S^ (Supplementary Fig. S12c). (Note that the *rpoS* gene in MG1655 is the pseudo revertant *rpoS* 33Q allele)(4, 12). Additionally, *E. coli* K-12 strains also carry changes in genes that influence translation. Like MG1655 and W3110, EMG2 carries a mutation in the gene encoding release factor RF2 (*prfB*) (Thr at position 246) and a mutation in *rpsG* (30S ribosomal protein S7), which results in C-terminal extension of the S7 protein product (Supplementary Fig. S12d and e). Both substitutions have been shown to affect translation, with the mutation in RF2 resulting in poor termination at UGA stop codons and the trans-translational tagging of S7 with the SsrA peptide (42-46). Thus, it is clear that, for both EMG2 and WG1, adaptation to a laboratory lifestyle has resulted in strains, with altered transcription and translation machineries, that likely impact on global gene expression.

Our analysis also indentifies mutations in genes involved in metabolism and cellular homeostasis (Supplementary File S1). Similar to MG1655 and W3110, EMG2 carries a frame shift in *rph* (previously termed *rph-1)* that results in a truncation of RNase PH, which affects the expression of *pyrE*, manifesting in a pyrimidine starvation phenotype (Supplementary Fig. S12f) (47, 48). Like other K-12 strains, EMG2 also carries a mutation in *ilvG*, which produces a truncated protein product that affects branch chained amino acid biosynthesis (49) (Supplementary Fig. S12g). Whilst these mutations are absent from WG1, WG1 carries lesions in *mdtF* (an AcrB efflux pump homologue) and *nfi* (DNA repair endonuclease V), both of which result in truncated products (Supplementary Fig. S11h and i). Thus, WG1 is likely compromised in both drug efflux and DNA damage repair (50, 51).

## DISCUSSION

The use of *E. coli* K-12 has shaped biological knowledge and research over the last century (52). Fred Neidhardt’s comment that ‘All cell biologists have at least two cells of interest: the one they are studying and *E. coli*’ (53) still holds true for many scientists, with *E. coli* K-12 still the cornerstone of molecular biology and microbiology. However, it is clear that adaptation to the laboratory lifestyle has resulted in *E. coli* K-12 strains which have alterations in transcription, translation, general metabolism and cellular homeostasis. As *E. coli* K-12 strains EMG2, MG1655 and W3110 share many common alterations (*e*.*g*., in *rpoD, prfB, rpsG, rph* (*rph-1*), *wbbL* (*rfb-50*), *prfB* and *ilvG*) this indicates that they share a similar lineage and that many of these mutations were fixed in their common ancestral strain (Fig. 1). On the other hand, WG1 carries alterations in different genes (e.g., *rfb-50, rpoA, mdtF* and *nfi*), suggesting that it is distinct from these strains (Fig. 1). It is worth noting that WG1 is similar to *E. coli* strains NCM3722 (54) and LS5218 (55). Strain NCM3722 (CGSC#12355) was first detailed by Sydney Kustu (48) and LS5218 is an industrial strain used for the production of fatty acid derived products (55). Both strains carry bacteriophage λ, a smaller version of the F plasmid (Tables 1 and 2: Supplementary Figs. S13 and S14) and contain many of the mutations carried by WG1 (54, 55).

In addition to lineage specific mutations, it is clear that WG1 and EMG2 have undergone their own lab-based evolution events, such as loss of cryptic prophages and gene disruption. The suggestion is that the selection of particular traits by microbiologists has driven lab-based evolution. Hence, IS inactivation of *finO* in F made plasmid transfer easier to study, larger plaques enabled the intricacies of λ lysogeny to be examined and lack of O-antigen enhances plasmid transformation (6, 7, 30, 35). Thus, our interpretation of *E. coli* biology has been inadvertently biased. Moreover, many other laboratory strains, handed down for generations, are as yet unsequenced, so it is unclear what other changes lie within those strains.

Heterogeneity in bacterial lab strains and plasmids has been observed many times and we are at a stage when even the same *E. coli* K-12 stock strains can produce different outcomes, calling reproducibility into question (27, 48, 56-59). It is clear that there are significant major differences between K-12 and other commensal *E. coli* strains, and these differences became fixed in the ancestors of the very widely used MG1655 and W3110 strains. Given the different mutations seen in WG1 compared to EMG2, it seems likely that identical or similar mutations will be present in other K-12 lineages. However, due to the extensive genetic systems that have been developed, demonstration of safe use, and lack of ability to colonize humans, *E. coli* K-12 strains will justifiably continue to be widely used (10, 11, 52). We think it is important that there is an awareness of the mutations present in K-12 strains, and the effects of these mutations on the physiology and metabolism of these strains. An understanding of the conditions that might select for mutants in laboratories, and the use of cost effective and accurate sequencing of laboratory stocks should help to prevent further undetected mutations arising in K-12 strains, which could compromise our understanding of fundamental biological processes. Thus, it is hoped that the next century will continue to provide more insight into the complex biology and evolution of this versatile organism. Indeed, appreciation of various K-12 strains, as well differences between various bacterial families, is sure to enhance our understanding of life.

## Abbreviations

ACT: Artemis Comparison Tool
BLAST: Basic Local Alignment Search Tool
CDS: coding sequence
CGE: Center for Genomic Epidemiology
CGSC: Coli Genetic Stock Centre
ONT: Oxford Nanopore Technologies
SNV: single nucleotide variant.

## Acknowledgements

We thank MicrobesNG for genome sequencing, in particular Andrew Holmes and Emily Jane Richardson for bioinformatics and SNV calling.

## Author contributions

DFB, JLH and SJWB conceived the study, selected samples, carried out bioinformatic analyses and wrote the manuscript. All authors read and approved the final version of the manuscript.

## Ethical statement

No ethical clearance was required for this study.

## Consent to publish

All authors give their consent to publish.

