## Supplementary Material for "Laboratory strains of *Escherichia coli* K-12: not such perfect role models after all"

**Edited by DFB on 26/06/22**

### Supplementary Material,

Douglas F. Browning [1]*, Jon L. Hobman [2], Stephen J.W. Busby[3].

[1] School of Biosciences, College of Health and Life Sciences, Aston University, Aston Triangle, Birmingham, B4 7ET, UK.

[2] School of Biosciences, University of Nottingham, Sutton Bonington Campus, Sutton Bonington, Loughborough, LE12 5RD, UK.

[3] Institute of Microbiology and Infection, School of Biosciences, University of Birmingham, Birmingham, B15 2TT, UK.

### Supplementary Figure legends

**Fig. S1.** Comparison of the genome of EMG2 with those of *E. coli* K-12 strains WG1, MG1655 and W3110. The figure shows the comparison of the EMG2 chromosome (contig 1) and F plasmid (contig 2) with the genomes from WG1, MG1655 (NC_000913.3) and W3110 (NC_007779.1), using the Proksee Server (https://proksee.ca/ ) (1). Selected features and regions of difference are labelled. The green, brown and blue rings illustrate the BLAST results when the genome sequences of *E. coli* K-12 strains WG1, MG1655 and W3110, respectively, are compared to that of EMG2.

**Fig. S2.** Chromosomal comparisons of different *E. coli* K-12 strains. The figure shows the comparison of the chromosomes from EMG2, WG1, MG1655 (NC_000913.3) and W3110 (NC_007779.1), using the Artemis Comparison Tool (ACT) (2).

**Fig. S3.** Comparison of the *E. coli* K-12 MG1655 chromosome with the genomes of *E. coli* K-12 strains WG1, EMG2 and W3110. The figure shows the comparison of the *E. coli* K-12 MG1655 chromosome (NC_000913.3) with those from WG1, EMG2 and W3110 (NC_007779.1), using the Proksee Server (1). The green, brown and blue rings illustrate the BLAST results when the genome sequences of *E. coli* K-12 strains WG1, EMG2 and W3110, respectively, are compared to that of MG1655.

**Fig. S4.** Detailed comparison of the *E. coli* K-12 MG1655 chromosome with other *E. coli* K-12 strains. The figure shows the comparison of regions of the *E. coli* K-12 MG1655 chromosome (NC_000913.3) with those from WG1, EMG2 and W3110 (NC_007779.1), using the Proksee Server (1). Panel a) shows the cryptic prophage CPZ-55 locus, b) the cryptic prophage CP4-6, c) the *rfb* cluster and d) the *gatYZABDR* locus.

**Fig. S5.** Comparison of the *gatYZABDR* locus from various *E. coli* K-12 strains. The figure shows the comparison of the *gatYZABDR* locus from EMG2, WG1, MG1655 (NC_000913.3) and W3110 (NC_007779.1), using (ACT) (2). Unique IS elements are labelled for MG1655 and W3110.

**Fig. S6.** Comparison of the *flhDC* region from various *E. coli* K-12 strains. The figure shows the comparison of the *E. coli* K-12 WG1 *flhDC* region with that of EMG2, MG1655 (NC_000913.3) and W3110 (NC_007779.1), using a) the Proksee Server (1) and b) ACT (2). In b) specific transposon and IS elements upstream of *flhDC* are indicated.

**Fig. S7.** Comparison of the region upstream of *yghO* in various *E. coli* K-12 strains. The figure shows the comparison of the region upstream of *yghO* in *E. coli* K-12 WG1 with that of EMG2, MG1655 (NC_000913.3) and W3110 (NC_007779.1), using a) the Proksee Server (1) and b) ACT (2). Genes encoding IS*5* transposases are highlighted. The genes *lptG*, *acpP* and *oleD* are predicted to encode an LPS export ABC transporter permease, an acyl-carrier protein and a NAD-dependent epimerase/dehydratase, respectively.

**Fig. S8.** Analysis of the F plasmid from WG1. The figure details the comparison of the F plasmid from WG1 with that from EMG2 and the previously sequenced F plasmid (AP001918.1) using Proksee [48]. The outer two rings display the genes and features of the WG1 F plasmid, with selected genes labelled. The green and brown rings illustrate the BLAST results when the F plasmid sequences of *E. coli* K-12 (AP001918.1) and EMG2, respectively, are compared to that from WG1.

**Fig. S9.** Comparison of F plasmids from different *E. coli* strains. a) The panel shows the comparison of the F plasmid (AP001918.1) with that from WG1 and EMG2, using ACT (2). Selected feature and regions of difference are labelled. b) Alignment of the F plasmid FinO proteins from different *E. coli* K-12 strains. The figure shows the alignment of the FinO proteins from WG1, EMG2 and the F plasmid (AP001918.1). The FinO EMG2 and F plasmid (AP001918.1) versions have been truncated by the insertion of an IS*3* insertion sequence.

**Fig. S10.** Comparison of phage λ from different *E. coli* strains. The figure shows the comparison of phage λ (NC_001416) with that from WG1 and EMG2, using ACT (2). Note that in WG1 and EMG2 λ is integrated into the bacterial chromosome as a prophage.

**Fig. S11.** Comparison of tail fibre proteins from phage λ. The figure shows the alignment of phage λ tail proteins, a) J, b) Stf and c) Tfa from the previous sequenced λ genome (NC_001416) with that from *E. coli* K-12 strains WG1 and EMG2. Differences are highlighted in bold and red.

**Fig. S12.** Comparison of various proteins that differ in *E. coli* K-12 genomes. The figure shows the alignment of proteins, a) RpoD (RNAP $\sigma^{70}$ subunit), b) RpoD (RNAP α subunit), c) RpoS (RNAP $\sigma^{S}$ subunit), d) PrfB (RF2), e) RpsG, f) Rph, g) IlvG, h) MdtF and i) Nfi from various *E. coli* strains, including WG1, EMG2, MG1655 (NC_000913.3), W3110 (NC_007779.1), EDL399 (NZ_CP008957), 042

(FN554766), BL21 (CP060121) and BW25113 (CP009273.1). Differences are highlighted in red.

**Fig. S13.** Comparison of the genome of WG1 with those of *E. coli* K-12 strains MG1655, NCM3722 and LS5218. The figure shows the comparison of the WG1 chromosome (contig 1) and F plasmid (contig 2) with the genomes from MG1655 (NC_000913.3), NCM3722 (CP011495.1 and CP011496.1) and LS5218 (MVJG00000000.1) using the Proksee Server (1). Selected features and regions of difference are labelled.

**Fig. S14.** Comparison of the F plasmid from *E. coli* K-12 strains EMG2, WG1, NCM3722 and LS5218. a) The panel shows the comparison of the F plasmid (AP001918.1) with that from EMG2, WG1, NCM3722 (CP011496.1) and LS5218 (MVJG00000000.1) using the Proksee Server (1). b) The panel shows the comparison of the F plasmid from WG1 and NCM3722 (CP011496.1), using ACT (2).

### References.


1. Grant JR, Stothard P. The CGView Server: a comparative genomics tool for circular genomes. Nucleic acids research. 2008;36(Web Server issue):W181-4.
2. Carver TJ, Rutherford KM, Berriman M, Rajandream MA, Barrell BG, Parkhill J. ACT: the Artemis Comparison Tool. Bioinformatics (Oxford, England). 2005;21(16):3422-3.

**Supplementary Fig. S1.**

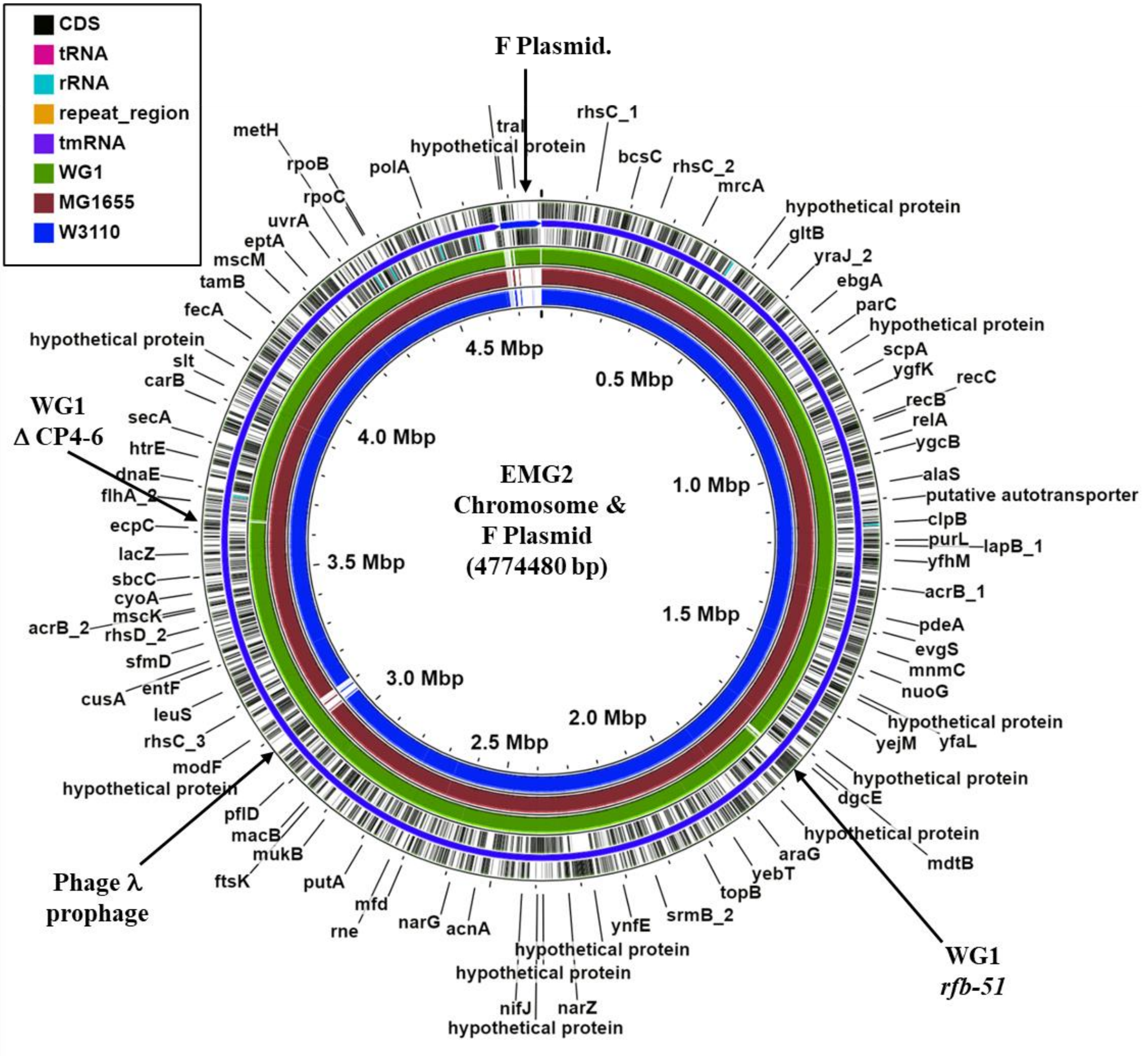

**Supplementary Fig. S2.**

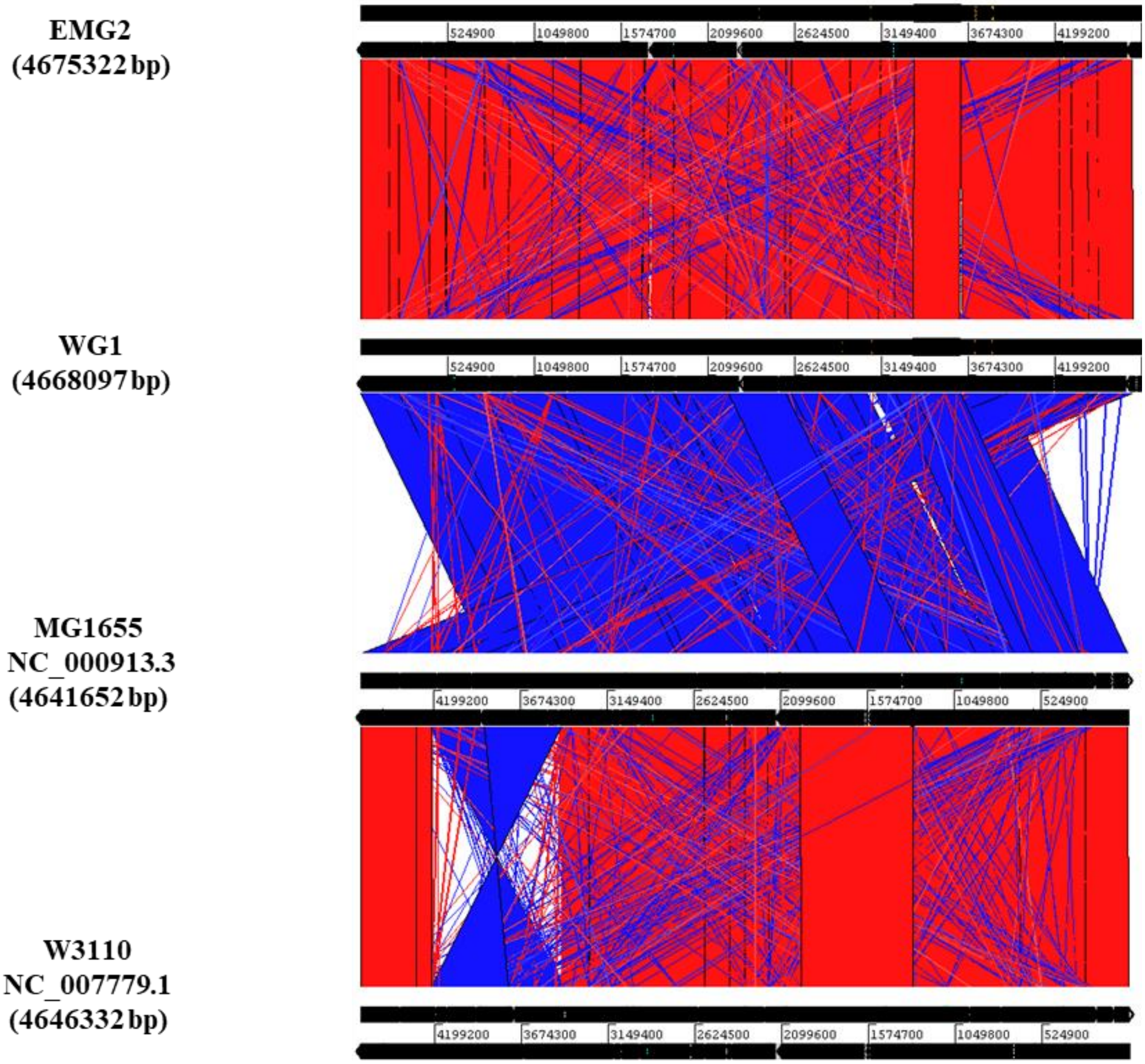

**Supplementary Fig. S3.**

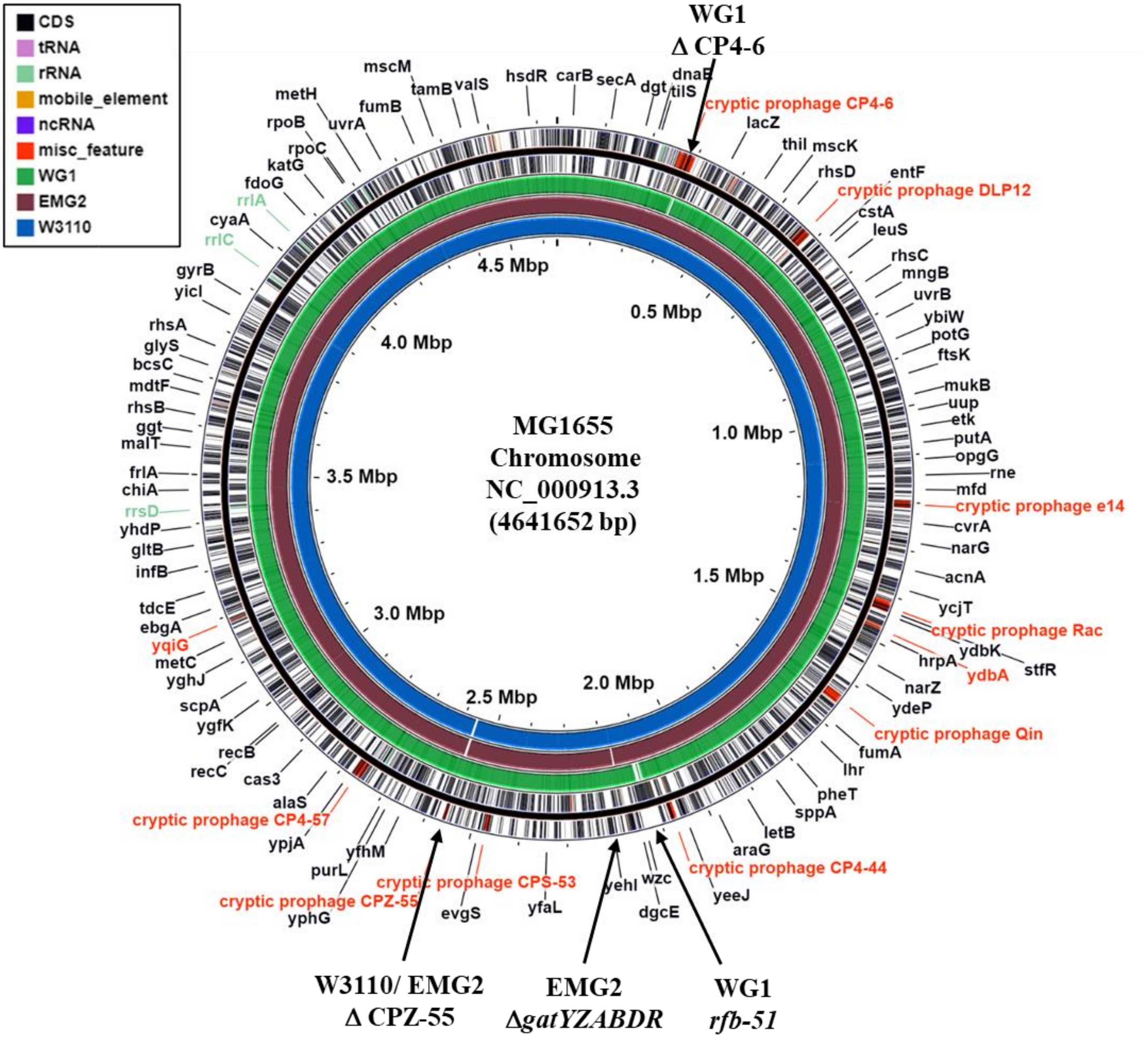

**Supplementary Fig. S4.**

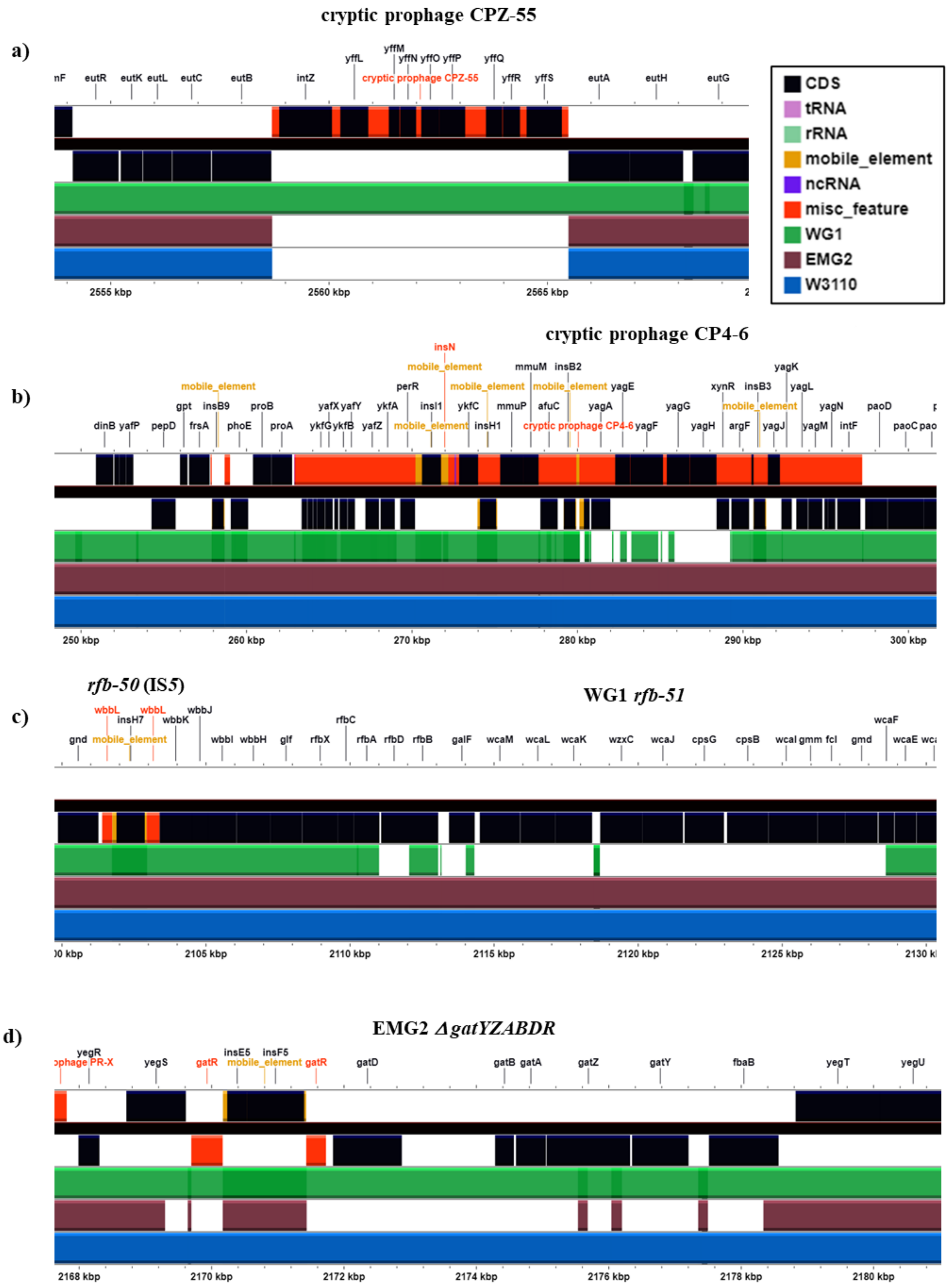

**Supplementary Fig. S5.**

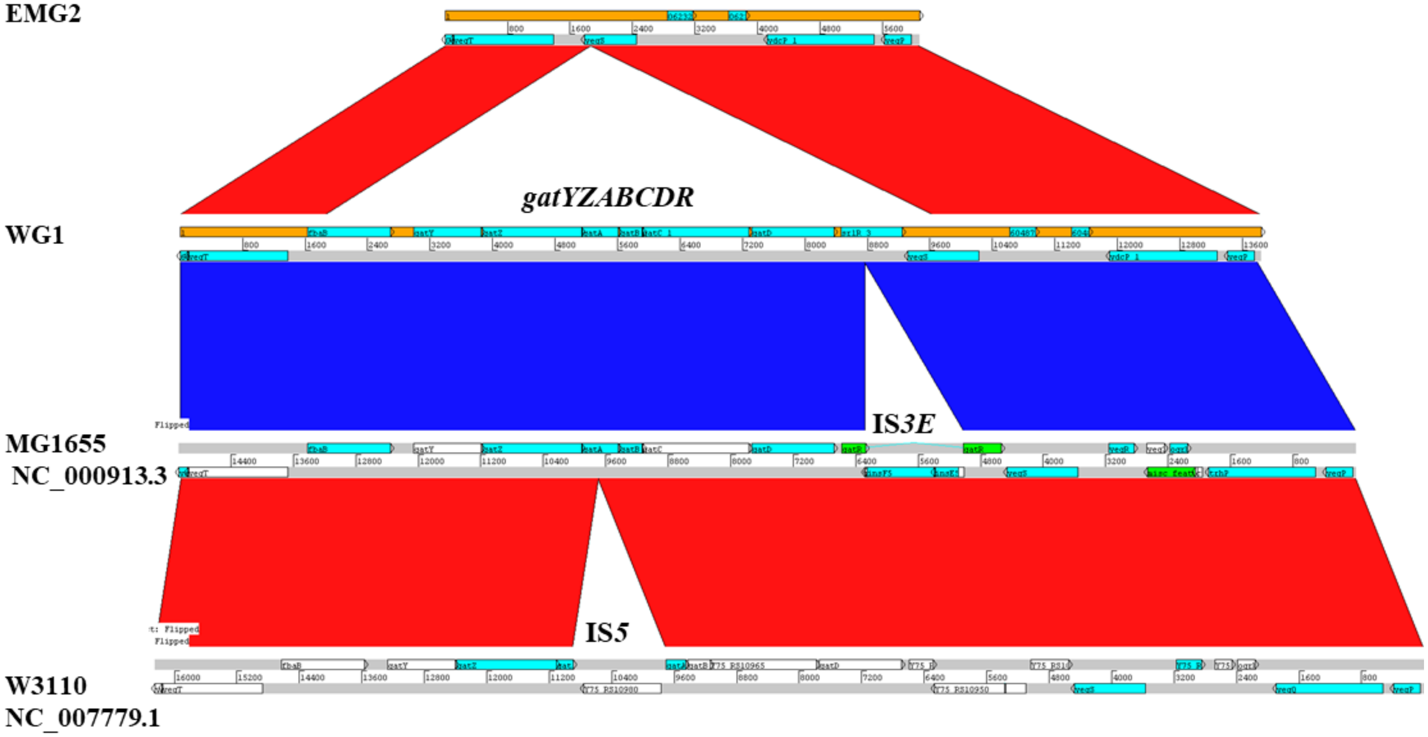

**Supplementary Fig. S6.**

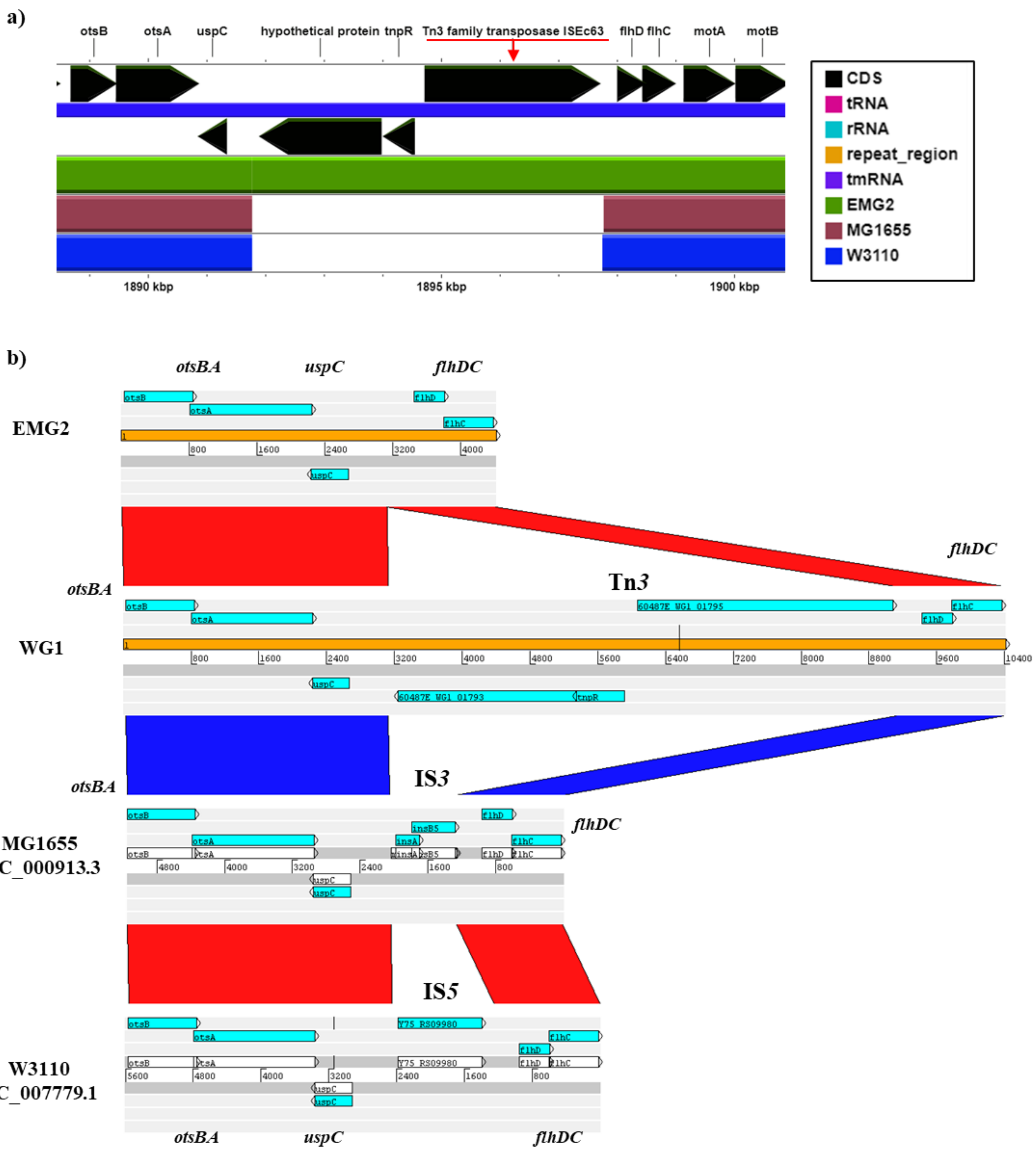

**Supplementary Fig. S7.**

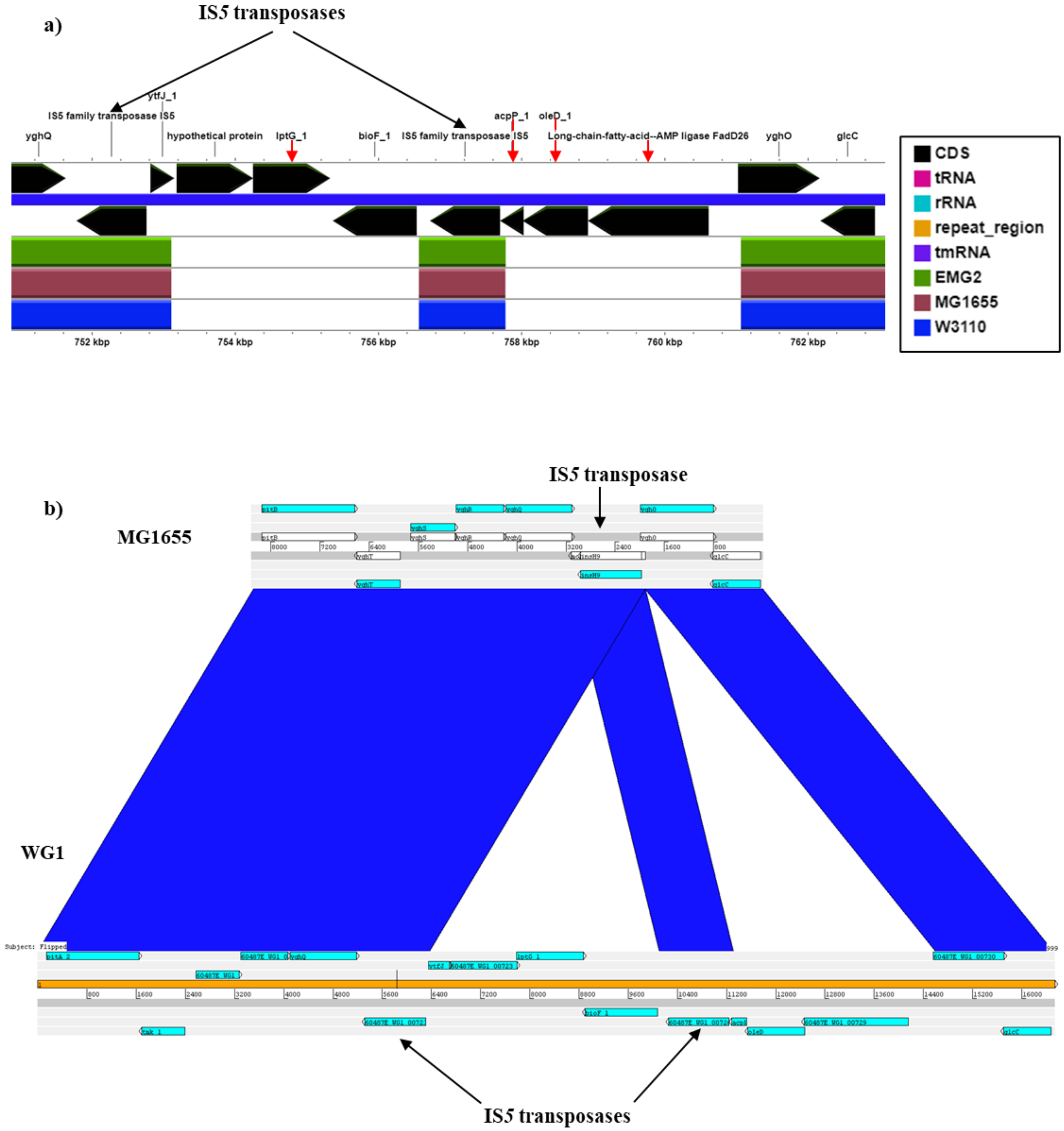

**Supplementary Fig. S8.**

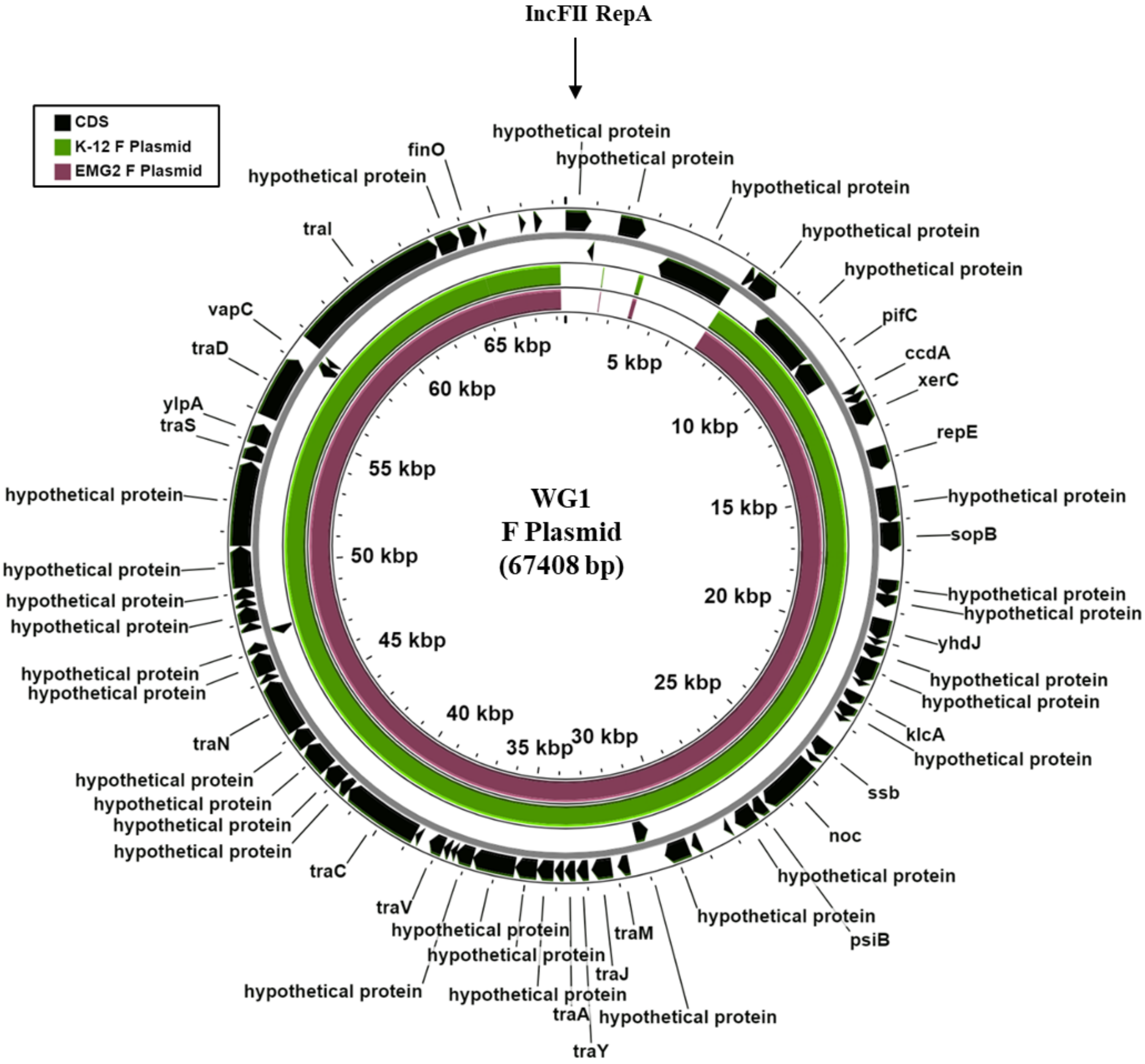

### Supplementary Fig. S9.

**a)**

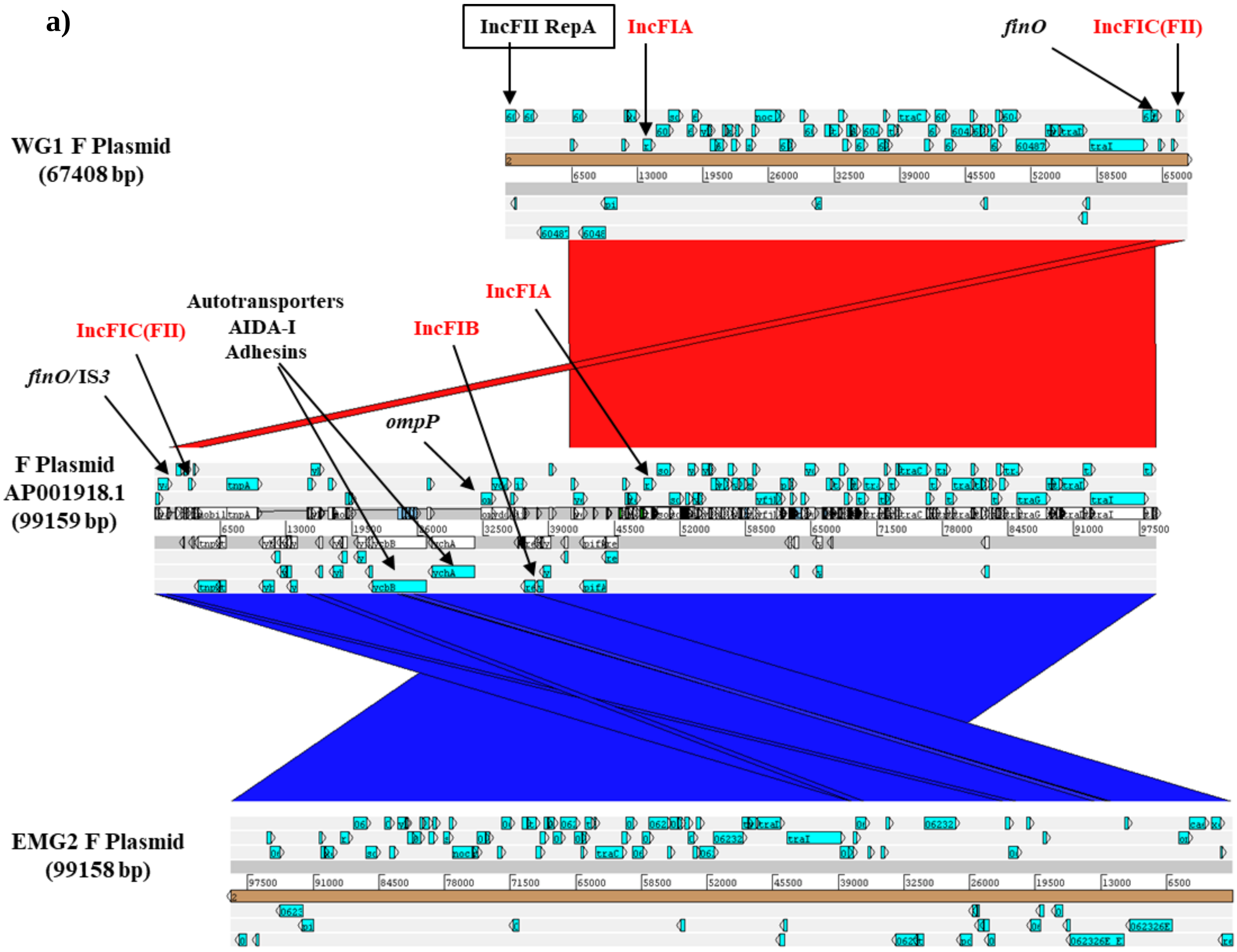


**b)**

```
FinO_WG1_       MTEQKRPVLTLKRKTEGETLVRSRKTIINVTTPPKWKVKKQKLAEKAAREAELAAKKAQA
FinO            MTEQKRPVLTLKRKTEGETLVRSRKTIINVTTPPKWKVKKQKLAEKAAREAELAAKKAQA
FinO_EMG2_      MTEQKRPVLTLKRKTEGETLVRSRKTIINVTTPPKWKVKKQKLAEKAAREAELAAKKAQA
                ************************************************************

FinO_WG1_       RQALSIYLNLPTLDDAVNTLKPWWPGLFDGDTPRLLACGIRDVLLEDVAQRNIPLSHKKL
FinO            RQALSIYLNLPTLDDAVNTLKPWWPGLFDGDTPRLLACGIRDVLLEDVAQRNIPLSHKKL
FinO_EMG2_      RQALSIYLNLPTLDDAVNTLKPWWPGLFDGDTPRLLACGIRDVLLEDVAQRNIPLSHKKL
                ************************************************************

FinO_WG1_       RRALKAITRSESYLCAMKAGACRYDTEGYVTEHISQEEEAYAAERLDKIRRQNRIKAELQ
FinO            RRALKAITRSE-------------------------------------------------
FinO_EMG2_      RRALKAITRSES------------------------------------------------
                ***********

FinO_WG1_       AVLDEK
FinO            ------
FinO_EMG2_      ------
```

**Supplementary Fig. S10.**

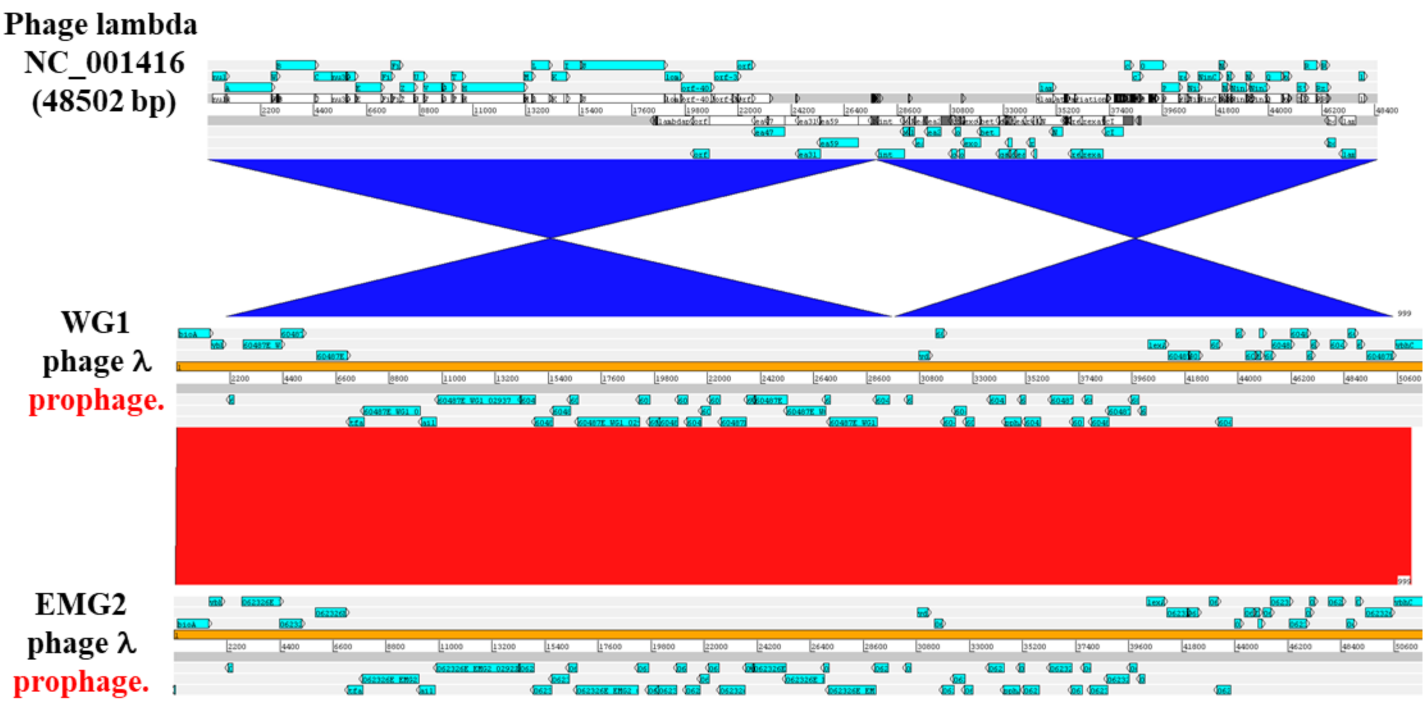

### Supplementary Fig. S11.

### a) Host specificity protein J (Tail tip protein).

```
J_WG1           MGKGSSKGHTPREAKDNLKSTQLLSVIDAISEGPIEGPVDGLKSVLLNSTPVLDTEGNTN
J_EMG2          MGKGSSKGHTPREAKDNLKSTQLLSVIDAISEGPIEGPVDGLKSVLLNSTPVLDTEGNTN
J               MGKGSSKGHTPREAKDNLKSTQLLSVIDAISEGPIEGPVDGLKSVLLNSTPVLDTEGNTN
                ************************************************************

J_WG1           ISGVTVVFRAGEQEQTPPEGFESSGSETVLGTEVKYDTPITRTITSANIDRLRFTFGVQA
J_EMG2          ISGVTVVFRAGEQEQTPPEGFESSGSETVLGTEVKYDTPITRTITSANIDRLRFTFGVQA
J               ISGVTVVFRAGEQEQTPPEGFESSGSETVLGTEVKYDTPITRTITSANIDRLRFTFGVQA
                ************************************************************

J_WG1           LVETTSKGDRNPSEVRLLVQIQRNGGWVTEKDITIKGKTTSQYLASVVMGNLPPRPFNIR
J_EMG2          LVETTSKGDRNPSEVRLLVQIQRNGGWVTEKDITIKGKTTSQYLASVVMGNLPPRPFNIR
J               LVETTSKGDRNPSEVRLLVQIQRNGGWVTEKDITIKGKTTSQYLASVVMGNLPPRPFNIR
                ************************************************************

J_WG1           MRRMTPDSTTDQLQNKTLWSSYTEIIDVKQCYPNTALVGVQVDSEQFGSQQVSRNYHLRG
J_EMG2          MRRMTPDSTTDQLQNKTLWSSYTEIIDVKQCYPNTALVGVQVDSEQFGSQQVSRNYHLRG
J               MRRMTPDSTTDQLQNKTLWSSYTEIIDVKQCYPNTALVGVQVDSEQFGSQQVSRNYHLRG
                ************************************************************

J_WG1           RILQVPSNYNPQTRQYSGIWDGTFKPAYSNNMAWCLWDMLTHPRYGMGKRLGAADVDKWA
J_EMG2          RILQVPSNYNPQTRQYSGIWDGTFKPAYSNNMAWCLWDMLTHPRYGMGKRLGAADVDKWA
J               RILQVPSNYNPQTRQYSGIWDGTFKPAYSNNMAWCLWDMLTHPRYGMGKRLGAADVDKWA
                ************************************************************

J_WG1           LYVIGQYCDQSVPDGFGGTEPRITCNAYLTTQRKAWDVLSDFCSAMRCMPVWNGQTLTFV
J_EMG2          LYVIGQYCDQSVPDGFGGTEPRITCNAYLTTQRKAWDVLSDFCSAMRCMPVWNGQTLTFV
J               LYVIGQYCDQSVPDGFGGTEPRITCNAYLTTQRKAWDVLSDFCSAMRCMPVWNGQTLTFV
                ************************************************************

J_WG1           QDRPSDKTWTYNRSNVVMPDDGAPFRYSFSALKDRHNAVEVNWIDPNNGWETATELVEDT
J_EMG2          QDRPSDKTWTYNRSNVVMPDDGAPFRYSFSALKDRHNAVEVNWIDPNNGWETATELVEDT
J               QDRPSDKTWTYNRSNVVMPDDGAPFRYSFSALKDRHNAVEVNWIDPNNGWETATELVEDT
                ************************************************************

J_WG1           QAIARYGRNVTKMDAFGCTSRGQAHRAGLWLIKTELLETQTVDFSVGAEGLRHVPGDVIE
J_EMG2          QAIARYGRNVTKMDAFGCTSRGQAHRAGLWLIKTELLETQTVDFSVGAEGLRHVPGDVIE
J               QAIARYGRNVTKMDAFGCTSRGQAHRAGLWLIKTELLETQTVDFSVGAEGLRHVPGDVIE
                ************************************************************

J_WG1           ICDDDYAGISTGGRVLAVNSQTRTLTLDREITLPSSGTALISLVDGSGNPVSVEVQSVTD
J_EMG2          ICDDDYAGISTGGRVLAVNSQTRTLTLDREITLPSSGTALISLVDGSGNPVSVEVQSVTD
J               ICDDDYAGISTGGRVLAVNSQTRTLTLDREITLPSSGTALISLVDGSGNPVSVEVQSVTD
                ************************************************************

J_WG1           GVKVKVSRVPDGVAEYSVWGLKLPTLRQRLFRCVSIRENDDGTYAITAVQHVPEKEAIVD
J_EMG2          GVKVKVSRVPDGVAEYSVWGLKLPTLRQRLFRCVSIRENDDGTYAITAVQHVPEKEAIVD
J               GVKVKVSRVPDGVAEYSVWELKLPTLRQRLFRCVSIRENDDGTYAITAVQHVPEKEAIVD
                ******************* ****************************************

J_WG1           NGAHFDGEQSGTVNGVTPPAVQHLTAEVTADSGEYQVLARWDTPKVVKGVSFLLRLTVTA
J_EMG2          NGAHFDGEQSGTVNGVTPPAVQHLTAEVTADSGEYQVLARWDTPKVVKGVSFLLRLTVTA
J               NGAHFDGEQSGTVNGVTPPAVQHLTAEVTADSGEYQVLARWDTPKVVKGVSFLLRLTVTA
                ************************************************************

J_WG1           DDGSERLVSTARTTETTYRFTQLALGNYRLTVRAVNAWGQQGDPASVSFRIAAPAAPSRI
J_EMG2          DDGSERLVSTARTTETTYRFTQLALGNYRLTVRAVNAWGQQGDPASVSFRIAAPAAPSRI
J               DDGSERLVSTARTTETTYRFTQLALGNYRLTVRAVNAWGQQGDPASVSFRIAAPAAPSRI
                ************************************************************

J_WG1           ELTPGYFQITATPHLAVYDPTVQFEFWFSEKQIADIRQVETSTRYLGTALYWIAASINIK
J_EMG2          ELTPGYFQITATPHLAVYDPTVQFEFWFSEKQIADIRQVETSTRYLGTALYWIAASINIK
J               ELTPGYFQITATPHLAVYDPTVQFEFWFSEKQIADIRQVETSTRYLGTALYWIAASINIK
                ************************************************************

J_WG1           PGHDYYFYIRSVNTVGKSAFVEAVGRASDDAEGYLDFFKGKITESHLGKELLEKVELTED
J_EMG2          PGHDYYFYIRSVNTVGKSAFVEAVGRASDDAEGYLDFFKGKITESHLGKELLEKVELTED
J               PGHDYYFYIRSVNTVGKSAFVEAVGRASDDAEGYLDFFKGKITESHLGKELLEKVELTED
                ************************************************************

J_WG1           NASRLEEFSKEWKDASDKWNAMWAVKIEQTKDGKHYVAGIGLSMEDTEEGKLSQFLVAAN
J_EMG2          NASRLEEFSKEWKDASDKWNAMWAVKIEQTKDGKHYVAGIGLSMEDTEEGKLSQFLVAAN
J               NASRLEEFSKEWKDASDKWNAMWAVKIEQTKDGKHYVAGIGLSMEDTEEGKLSQFLVAAN
                ************************************************************
```

```
J_WG1           RIAFIDPANGNETPMFVAQGNQIFMNDVFLKRLTAPTITSGGNPPAFSLTPDGKLTAKNA
J_EMG2          RIAFIDPANGNETPMFVAQGNQIFMNDVFLKRLTAPTITSGGNPPAFSLTPDGKLTAKNA
J               RIAFIDPANGNETPMFVAQGNQIFMNDVFLKRLTAPTITSGGNPPAFSLTPDGKLTAKNA
                ************************************************************

J_WG1           DISGSVNANSGTLSNVTIAENCTINGTLRAEKIVGDIVKAASAAFPRQRESSVDWPSGTR
J_EMG2          DISGSVNANSGTLSNVTIAENCTINGTLRAEKIVGDIVKAASAAFPRQRESSVDWPSGTR
J               DISGSVNANSGTLSNVTIAENCTINGTLRAEKIVGDIVKAASAAFPRQRESSVDWPSGTR
                ************************************************************

J_WG1           TVTVTDDHPFDRQIVVLPLTFRGSKRTVSGRTTYSMCYLKVLMNGAVIYDGAANEAVQVF
J_EMG2          TVTVTDDHPFDRQIVVLPLTFRGSKRTVSGRTTYSMCYLKVLMNGAVIYDGAANEAVQVF
J               TVTVTDDHPFDRQIVVLPLTFRGSKRTVSGRTTYSMCYLKVLMNGAVIYDGAANEAVQVF
                ************************************************************

J_WG1           SRIVDMPAGRGNVILTFTLTSTRHSADIPPDTFASDVQVMVIKKQALGISVV
J_EMG2          SRIVDMPAGRGNVILTFTLTSTRHSADIPPDTFASDVQVMVIKKQALGISVV
J               SRIVDMPAGRGNVILTFTLTSTRHSADIPPYTFASDVQVMVIKKQALGISVV
                ****************************** *********************
```

### b) Stf (Side tail fibre).

```
stf_WG1          MAVKISGVLKDGTGKPVQNCTIQLKARRNSTTVVVNTVGSENPDEAGRYSMDVEYGQYSV
stf_orf-401      MAVKISGVLKDGTGKPVQNCTIQLKARRNSTTVVVNTVGSENPDEAGRYSMDVEYGQYSV
stf_EMG2         MAVKISGVLKDGTGKPVQNCTIQLKARRNSTTVVVNTVGSENPDEAGRYSMDVEYGQYSV
                 ************************************************************

stf_WG1          ILQVDGFPPSHAGTITVYEDSQPGTLNDFLCAMTEDDARPEVLRRLELMVEEVARNASVV
stf_orf-401      ILQVDGFPPSHAGTITVYEDSQPGTLNDFLCAMTEDDARPEVLRRLELMVEEVARNASVV
stf_EMG2         ILQVDGFPPSHAGTITVYEDSQPGTLNDFLCAMTEDDARPEVLRRLELMVEEVARNASVV
                 ************************************************************

stf_WG1          AQSTADAKKSAGDASASAAQVAALVTDATDSARAASTSAGQAASSAQEASSGAEAASAKA
stf_orf-401      AQSTADAKKSAGDASASAAQVAALVTDATDSARAASTSAGQAASSAQEASSGAEAASAKA
stf_EMG2         AQSTADAKKSAGDASASAAQVAALVTDATDSARAASTSAGQAASSAQEASSGAEAASAKA
                 ************************************************************

stf_WG1          TEAEKSAAAAESSKNAAATSAGAAKTSETNAAASQQSAATSASTAATKASEAATSARDAV
stf_orf-401      TEAEKSAAAAESSKNAAATSAGAAKTSETNAAASQQSAATSASTAATKASEAATSARDAV
stf_EMG2         TEAEKSAAAAESSKNAAATSAGAAKTSETNAAASQQSAATSASTAATKASEAATSARDAV
                 ************************************************************

stf_WG1          ASKEAAKSSETNASSSAGRAASSATAAENSARAAKTSETNARSSETAAERSASAAADAKT
stf_orf-401      ASKEAAKSSETNASSSAGRAASSATAAENSARAAKTSETNARSSETAAERSASAAADAKT
stf_EMG2         ASKEAAKSSETNASSSAGRAASSATAAENSARAAKTSETNARSSETAAERSASAAADAKT
                 ************************************************************

stf_WG1          AAAGSASTASTKATEAAGSAVSASQSKSAAEAAAIRAENSAKRAEDIASAVALEDADTTR
stf_orf-401      AAAGSASTASTKATEAAGSAVSASQSKSAAEAAAIRAKNSAKRAEDIASAVALEDADTTR
stf_EMG2         AAAGSASTASTKATEAAGSAVSASQSKSAAEAAAIRAENSAKRAEDIASAVALEDADTTR
                 *************************************:**********************

stf_WG1          KGIVQLSSATNSTSETLAATPKAVKVVMDETNRKAPLDSPALTGTPTAPTALRGTNNTQI
stf_orf-401      KGIVQLSSATNSTSETLAATPKAVKVVMDETNRKAHWTVRH-------------------
stf_EMG2         KGIVQLSSATNSTSETLAATPKAVKVVMDETNRKAPLDSPALTGTPTAPTALRGTNNTQI
                 ***********************************

stf_WG1          ANTAFVLAAIADVIDASPDALNTLNELAAALGNDPDFATTMTNALAGKQPKNATLTALAG
stf_orf-401      ------------------------------------------------------------
stf_EMG2         ANTAFVLAAIADVIDASPDALNTLNELAAALGNDPDFATTMTNALAGKQPKNATLTALAG

stf_WG1          LSTAKNKLPYFAENDAASLTELTQVGRDILAKNSVADVLEYLGAGENSAFPAGAPIPWPS
stf_orf-401      ------------------------------------------------------------
stf_EMG2         LSTAKNKLPYFAENDAASLTELTQVGRDILAKNSVADVLEYLGAGENSAFPAGAPIPWPS

stf_WG1          DIVPSGYVLMQGQAFDKSAYPKLAVAYPSGVLPDMRGWTIKGKPASGRAVLSQEQDGIKS
stf_orf-401      ------------------------------------------------------------
stf_EMG2         DIVPSGYVLMQGQAFDKSAYPKLAVAYPSGVLPDMRGWTIKGKPASGRAVLSQEQDGIKS

stf_WG1          HTHSASASGTDLGTKTTSSFDYGTKTTGSFDYGTKSTNNTGAHAHSLSGSTGAAGAHAHT
stf_orf-401      ------------------------------------------------------------
stf_EMG2         HTHSASASGTDLGTKTTSSFDYGTKTTGSFDYGTKSTNNTGAHAHSLSGSTGAAGAHAHT
```

```
stf_WG1          SGLRMNSSGWSQYGTATITGSLSTVKGTNTQGIAYLSKTDSQGSHSHSLSGTAVSAGAHA
stf_orf-401      ------------------------------------------------------------
stf_EMG2         SGLRMNSSGWSQYGTATITGSLSTVKGTNTQGIAYLSKTDSQGSHSHSLSGTAVSAGAHA


stf_WG1          HTVGIGAHQHPVVIGAHAHSFSIGSHGHTITVNAAGNAENTVKNIAFNYIVRLA
stf_orf-401      ------------------------------------------------------
stf_EMG2         HTVGIGAHQHPVVIGAHAHSFSIGSHGHTITVNAAGNAENTVKNIAFNYIVRLA
```

### c) Tra (Tail fibre addition).

```
TfaE_5_EMG2      MAFRMSEQPRTIKIYNLLAGTNEFIGEGDAYIPPHTGLPANSTDIAPPDIPAGFVAVFNS
TfaE_5_WG1       MAFRMSEQPRTIKIYNLLAGTNEFIGEGDAYIPPHTGLPANSTDIAPPDIPAGFVAVFNS
Tfa_orf-194      MAFRMSEQPRTIKIYNLLAGTNEFIGEGDAYIPPHTGLPANSTDIAPPDIPAGFVAVFNS
                 ************************************************************

TfaE_5_EMG2      DEASWHLVEDHRGKTVYDVASGDALFISELGPLPENFTWLSPGGEYQKWNGTAWVKDTEA
TfaE_5_WG1       DEASWHLVEDHRGKTVYDVASGDALFISELGPLPENFTWLSPGGEYQKWNGTAWVKDTEA
Tfa_orf-194      DEASWHLVEDHRGKTVYDVASGDALFISELGPLPENFTWLSPGGEYQKWNGTAWVKDTEA
                 ************************************************************

TfaE_5_EMG2      EKLFRIREAEETKKSLMQVASEHIAPLQDAADLEIATEEETSLLEAWKKYRVLLNRVDTS
TfaE_5_WG1       EKLFRIREAEETKKSLMQVASEHIAPLQDAADLEIATEEETSLLEAWKKYRVLLNRVDTS
Tfa_orf-194      EKLFRIREAEETKKSLMQVASEHIAPLQDAADLEIATKEETSLLEAWKKYRVLLNRVDTS
                 *************************************:**********************

TfaE_5_EMG2      TAPDIEWPAVPVME
TfaE_5_WG1       TAPDIEWPAVPVME
Tfa_orf-194      TAPDIEWPAVPVME
                 **************
```

**Supplementary Fig. S12.**

**a) $\sigma^{70}$ (RpoD)**

```
RpoD_WG1         MEQNPQSQLKLLVTRGKEQGYLTYAEVNDHLPEDIVDSDQIEDIIQMINDMGIQVMEEAP
RpoD_Bl21        MEQNPQSQLKLLVTRGKEQGYLTYAEVNDHLPEDIVDSDQIEDIIQMINDMGIQVMEEAP
RpoD_EMG2        MEQNPQSQLKLLVTRGKEQGYLTYAEVNDHLPEDIVDSDQIEDIIQMINDMGIQVMEEAP
RpoD_MG1655      MEQNPQSQLKLLVTRGKEQGYLTYAEVNDHLPEDIVDSDQIEDIIQMINDMGIQVMEEAP
RpoD_W3110       MEQNPQSQLKLLVTRGKEQGYLTYAEVNDHLPEDIVDSDQIEDIIQMINDMGIQVMEEAP
RpoD_BW25113     MEQNPQSQLKLLVTRGKEQGYLTYAEVNDHLPEDIVDSDQIEDIIQMINDMGIQVMEEAP
RpoD_042         MEQNPQSQLKLLVTRGKEQGYLTYAEVNDHLPEDIVDSDQIEDIIQMINDMGIQVMEEAP
RpoD_EDL399      MEQNPQSQLKLLVTRGKEQGYLTYAEVNDHLPEDIVDSDQIEDIIQMINDMGIQVMEEAP
                 ************************************************************

RpoD_WG1         DADDLMLAENTADEDAAEAAQVLSSVESEIGRTTDPVRMYMREMGTVELLTREGEIDIA
RpoD_Bl21        DADDLMLAENTADEDAAEAAQVLSSVESEIGRTTDPVRMYMREMGTVELLTREGEIDIA
RpoD_EMG2        DADDLMLAENTADEDAAEAAQVLSSVESEIGRTTDPVRMYMREMGTVELLTREGEIDIA
RpoD_MG1655      DADDLMLAENTADEDAAEAAQVLSSVESEIGRTTDPVRMYMREMGTVELLTREGEIDIA
RpoD_W3110       DADDLMLAENTADEDAAEAAQVLSSVESEIGRTTDPVRMYMREMGTVELLTREGEIDIA
RpoD_BW25113     DADDLMLAENTADEDAAEAAQVLSSVESEIGRTTDPVRMYMREMGTVELLTREGEIDIA
RpoD_042         DADDLMLAENTADEDAAEAAQVLSSVESEIGRTTDPVRMYMREMGTVELLTREGEIDIA
RpoD_EDL399      DADDLMLAENTADEDAAEAAQVLSSVESEIGRTTDPVRMYMREMGTVELLTREGEIDIA
                 ***********************************************************

RpoD_WG1         KRIEDGINQVQCSVAEYPEAITYLLEQYDRVEAEEARLSDLITGFVDPNAEEDLAPTATH
RpoD_Bl21        KRIEDGINQVQCSVAEYPEAITYLLEQYDRVEAEEARLSDLITGFVDPNAEEDLAPTATH
RpoD_EMG2        KRIEDGINQVQCSVAEYPEAITYLLEQYDRVEAEEARLSDLITGFVDPNAEEDLAPTATH
RpoD_MG1655      KRIEDGINQVQCSVAEYPEAITYLLEQYDRVEAEEARLSDLITGFVDPNAEEDLAPTATH
RpoD_W3110       KRIEDGINQVQCSVAEYPEAITYLLEQYDRVEAEEARLSDLITGFVDPNAEEDLAPTATH
RpoD_BW25113     KRIEDGINQVQCSVAEYPEAITYLLEQYDRVEAEEARLSDLITGFVDPNAEEDLAPTATH
RpoD_042         KRIEDGINQVQCSVAEYPEAITYLLEQYDRVEAEEARLSDLITGFVDPNAEEDLAPTATH
RpoD_EDL399      KRIEDGINQVQCSVAEYPEAITYLLEQYDRVEAEEARLSDLITGFVDPNAEEDLAPTATH
                 ************************************************************

RpoD_WG1         VGSELSQEDLDDDEDEDEEDGDDDSADDDNSIDPELAREKFAELRAQYVVTRDTIKAKGR
RpoD_Bl21        VGSELSQEDLDDDEDEDEEDGDDDSADDDNSIDPELAREKFAELRAQYVVTRDTIKAKGR
RpoD_EMG2        VGSELSQEDLDDDEDEDEEDGDDDSADDDNSIDPELAREKFAELRAQYVVTRDTIKAKGR
RpoD_MG1655      VGSELSQEDLDDDEDEDEEDGDDDSADDDNSIDPELAREKFAELRAQYVVTRDTIKAKGR
RpoD_W3110       VGSELSQEDLDDDEDEDEEDGDDDSADDDNSIDPELAREKFAELRAQYVVTRDTIKAKGR
RpoD_BW25113     VGSELSQEDLDDDEDEDEEDGDDDSADDDNSIDPELAREKFAELRAQYVVTRDTIKAKGR
RpoD_042         VGSELSQEDLDDDEDEDEEDGDDDSADDDNSIDPELAREKFAELRAQYVVTRDTIKAKGR
RpoD_EDL399      VGSELSQEDLDDDEDEDEEDGDDDSADDDNSIDPELAREKFAELRAQYVVTRDTIKAKGR
                 ************************************************************

RpoD_WG1         SHATAQEEILKLSEVFKQFRLVPKQFDYLVNSMRVMMDRVRTQERLIMKLCVEQCKMPKK
RpoD_Bl21        SHATAQEEILKLSEVFKQFRLVPKQFDYLVNSMRVMMDRVRTQERLIMKLCVEQCKMPKK
RpoD_EMG2        SHATAQEEILKLSEVFKQFRLVPKQFDYLVNSMRVMMDRVRTQERLIMKLCVEQCKMPKK
RpoD_MG1655      SHATAQEEILKLSEVFKQFRLVPKQFDYLVNSMRVMMDRVRTQERLIMKLCVEQCKMPKK
RpoD_W3110       SHATAQEEILKLSEVFKQFRLVPKQFDYLVNSMRVMMDRVRTQERLIMKLCVEQCKMPKK
RpoD_BW25113     SHATAQEEILKLSEVFKQFRLVPKQFDYLVNSMRVMMDRVRTQERLIMKLCVEQCKMPKK
RpoD_042         SHAAAQEEILKLSEVFKQFRLVPKQFDYLVNSMRVMMDRVRTQERLIMKLCVEQCKMPKK
RpoD_EDL399      SHAAAQEEILKLSEVFKQFRLVPKQFDYLVNSMRVMMDRVRTQERLIMKLCVEQCKMPKK
                 ***:********************************************************

RpoD_WG1         NFITLFTGNETSDTWFNAAIAMNKPWSEKLHDVSEEVHRALQKLQQIEEETGLTIEQVKD
RpoD_Bl21        NFITLFTGNETSDTWFNAAIAMNKPWSEKLHDVSEEVHRALQKLQQIEEETGLTIEQVKD
RpoD_EMG2        NFITLFTGNETSDTWFNAAIAMNKPWSEKLHDVSEEVHRALQKLQQIEEETGLTIEQVKD
RpoD_MG1655      NFITLFTGNETSDTWFNAAIAMNKPWSEKLHDVSEEVHRALQKLQQIEEETGLTIEQVKD
RpoD_W3110       NFITLFTGNETSDTWFNAAIAMNKPWSEKLHDVSEEVHRALQKLQQIEEETGLTIEQVKD
RpoD_BW25113     NFITLFTGNETSDTWFNAAIAMNKPWSEKLHDVSEEVHRALQKLQQIEEETGLTIEQVKD
RpoD_042         NFITLFTGNETSDTWFNAAIAMNKPWSEKLHDVSEEVHRALQKLQQIEEETGLTIEQVKD
RpoD_EDL399      NFITLFTGNETSDTWFNAAIAMNKPWSEKLHDVSEEVHRALQKLQQIEEETGLTIEQVKD
                 ************************************************************
```

```
RpoD_WG1          INRRMSIGEAKARRAKKEMVEANLRLVISIAKKYTNRGLQFLDLIQEGNIGLMKAVDKFE
RpoD_Bl21         INRRMSIGEAKARRAKKEMVEANLRLVISIAKKYTNRGLQFLDLIQEGNIGLMKAVDKFE
RpoD_EMG2         INRRMSIGEAKARRAKKEMVEANLRLVISIAKKYTNRGLQFLDLIQEGNIGLMKAVDKFE
RpoD_MG1655       INRRMSIGEAKARRAKKEMVEANLRLVISIAKKYTNRGLQFLDLIQEGNIGLMKAVDKFE
RpoD_W3110        INRRMSIGEAKARRAKKEMVEANLRLVISIAKKYTNRGLQFLDLIQEGNIGLMKAVDKFE
RpoD_BW25113      INRRMSIGEAKARRAKKEMVEANLRLVISIAKKYTNRGLQFLDLIQEGNIGLMKAVDKFE
RpoD_042          INRRMSIGEAKARRAKKEMVEANLRLVISIAKKYTNRGLQFLDLIQEGNIGLMKAVDKFE
RpoD_EDL399       INRRMSIGEAKARRAKKEMVEANLRLVISIAKKYTNRGLQFLDLIQEGNIGLMKAVDKFE
                  ************************************************************

RpoD_WG1          YRRGYKFSTYATWWIRQAITRSIADQARTIRIPVHMIETINKLNRISRQMLQEMGREPTP
RpoD_Bl21         YRRGYKFSTYATWWIRQAITRSIADQARTIRIPVHMIETINKLNRISRQMLQEMGREPTP
RpoD_EMG2         YRRGYKFSTYATWWIRQAITRSIADQARTIRIPVHMIETINKLNRISRQMLQEMGREPTP
RpoD_MG1655       YRRGYKFSTYATWWIRQAITRSIADQARTIRIPVHMIETINKLNRISRQMLQEMGREPTP
RpoD_W3110        YRRGYKFSTYATWWIRQAITRSIADQARTIRIPVHMIETINKLNRISRQMLQEMGREPTP
RpoD_BW25113      YRRGYKFSTYATWWIRQAITRSIADQARTIRIPVHMIETINKLNRISRQMLQEMGREPTP
RpoD_042          YRRGYKFSTYATWWIRQAITRSIADQARTIRIPVHMIETINKLNRISRQMLQEMGREPTP
RpoD_EDL399       YRRGYKFSTYATWWIRQAITRSIADQARTIRIPVHMIETINKLNRISRQMLQEMGREPTP
                  ************************************************************

RpoD_WG1          EELAERMLMPEDKIRKVLKIAKEPISMETPIGDDEDSHLGDFIEDTTLELPLDSATTESL
RpoD_Bl21         EELAERMLMPEDKIRKVLKIAKEPISMETPIGDDEDSHLGDFIEDTTLELPLDSATTESL
RpoD_EMG2         EELAERMLMPEDKIRKVLKIAKEPISMETPIGDDEDSHLGDFIEDTTLELPLDSATTESL
RpoD_MG1655       EELAERMLMPEDKIRKVLKIAKEPISMETPIGDDEDSHLGDFIEDTTLELPLDSATTESL
RpoD_W3110        EELAERMLMPEDKIRKVLKIAKEPISMETPIGDDEDSHLGDFIEDTTLELPLDSATTESL
RpoD_BW25113      EELAERMLMPEDKIRKVLKIAKEPISMETPIGDDEDSHLGDFIEDTTLELPLDSATTESL
RpoD_042          EELAERMLMPEDKIRKVLKIAKEPISMETPIGDDEDSHLGDFIEDTTLELPLDSATTESL
RpoD_EDL399       EELAERMLMPEDKIRKVLKIAKEPISMETPIGDDEDSHLGDFIEDTTLELPLDSATTESL
                  ************************************************************

                                              571  Helix turn Helix
RpoD_WG1          RAATHDVLAGLTAREAKVLRMRFGIDMNTDHTLEEVGKQFDVTRERIRQIEAKALRKLRH
RpoD_Bl21         RAATHDVLAGLTAREAKVLRMRFGIDMNTDHTLEEVGKQFDVTRERIRQIEAKALRKLRH
RpoD_EMG2         RAATHDVLAGLTAREAKVLRMRFGIDMNTDYTLEEVGKQFDVTRERIRQIEAKALRKLRH
RpoD_MG1655       RAATHDVLAGLTAREAKVLRMRFGIDMNTDYTLEEVGKQFDVTRERIRQIEAKALRKLRH
RpoD_W3110        RAATHDVLAGLTAREAKVLRMRFGIDMNTDYTLEEVGKQFDVTRERIRQIEAKALRKLRH
RpoD_BW25113      RAATHDVLAGLTAREAKVLRMRFGIDMNTDYTLEEVGKQFDVTRERIRQIEAKALRKLRH
RpoD_042          RAATHDVLAGLTAREAKVLRMRFGIDMNTDHTLEEVGKQFDVTRERIRQIEAKALRKLRH
RpoD_EDL399       RAATHDVLAGLTAREAKVLRMRFGIDMNTDHTLEEVGKQFDVTRERIRQIEAKALRKLRH
                  ******************************:*****************************

RpoD_WG1          PSRSEVLRSFLDD
RpoD_Bl21         PSRSEVLRSFLDD
RpoD_EMG2         PSRSEVLRSFLDD
RpoD_MG1655       PSRSEVLRSFLDD
RpoD_W3110        PSRSEVLRSFLDD
RpoD_BW25113      PSRSEVLRSFLDD
RpoD_042          PSRSEVLRSFLDD
RpoD_EDL399       PSRSEVLRSFLDD
                  *************
```

### b) α (RpoA)

```
RpoA_EDL399      MQGSVTEFLKPRLVDIEQVSSTHAKVTLEPLERGFGHTLGNALRRILLSSMPGCAVTEVE
RpoA_W3110       MQGSVTEFLKPRLVDIEQVSSTHAKVTLEPLERGFGHTLGNALRRILLSSMPGCAVTEVE
RpoA_042         MQGSVTEFLKPRLVDIEQVSSTHAKVTLEPLERGFGHTLGNALRRILLSSMPGCAVTEVE
RpoA_MG1655      MQGSVTEFLKPRLVDIEQVSSTHAKVTLEPLERGFGHTLGNALRRILLSSMPGCAVTEVE
RpoA_EMG2        MQGSVTEFLKPRLVDIEQVSSTHAKVTLEPLERGFGHTLGNALRRILLSSMPGCAVTEVE
RpoA_WG1         MQGSVTEFLKPRLVDIEQVSSTHAKVTLEPLERGFGHTLGNALRRILLSSMPGCAVTEVE
                 ************************************************************

RpoA_EDL399      IDGVLHEYSTKEGVQEDILEILLNLKGLAVRVQGKDEVILTLNKSGIGPVTAADITHDGD
RpoA_W3110       IDGVLHEYSTKEGVQEDILEILLNLKGLAVRVQGKDEVILTLNKSGIGPVTAADITHDGD
RpoA_042         IDGVLHEYSTKEGVQEDILEILLNLKGLAVRVQGKDEVILTLNKSGIGPVTAADITHDGD
RpoA_MG1655      IDGVLHEYSTKEGVQEDILEILLNLKGLAVRVQGKDEVILTLNKSGIGPVTAADITHDGD
RpoA_EMG2        IDGVLHEYSTKEGVQEDILEILLNLKGLAVRVQGKDEVILTLNKSGIGPVTAADITHDGD
RpoA_WG1         IDGVLHEYSTKEGVQEDILEILLNLKGLAVRVQGKDEVILTLNKSGIGPVTAADITHDGD
                 ************************************************************

RpoA_EDL399      VEIVKPQHVICHLTDENASISMRIKVQRGRGYVPASTRIHSEEDERPIGRLLVDACYSPV
RpoA_W3110       VEIVKPQHVICHLTDENASISMRIKVQRGRGYVPASTRIHSEEDERPIGRLLVDACYSPV
RpoA_042         VEIVKPQHVICHLTDENASISMRIKVQRGRGYVPASTRIHSEEDERPIGRLLVDACYSPV
RpoA_MG1655      VEIVKPQHVICHLTDENASISMRIKVQRGRGYVPASTRIHSEEDERPIGRLLVDACYSPV
RpoA_EMG2        VEIVKPQHVICHLTDENASISMRIKVQRGRGYVPASTRIHSEEDERPIGRLLVDACYSPV
RpoA_WG1         VEIVKPQHVICHLTDENASISMRIKVQRGRGYVPASTRIHSEEDERPIGRLLVDACYSPV
                 ************************************************************

RpoA_EDL399      ERIAYNVEAARVEQRTDLDKLVIEMETNGTIDPEEAIRRAATILAEQLEAFVDLRDVRQP
RpoA_W3110       ERIAYNVEAARVEQRTDLDKLVIEMETNGTIDPEEAIRRAATILAEQLEAFVDLRDVRQP
RpoA_042         ERIAYNVEAARVEQRTDLDKLVIEMETNGTIDPEEAIRRAATILAEQLEAFVDLRDVRQP
RpoA_MG1655      ERIAYNVEAARVEQRTDLDKLVIEMETNGTIDPEEAIRRAATILAEQLEAFVDLRDVRQP
RpoA_EMG2        ERIAYNVEAARVEQRTDLDKLVIEMETNGTIDPEEAIRRAATILAEQLEAFVDLRDVRQP
RpoA_WG1         ERIAYNVEAARVEQRTDLDKLVIEMETNGTIDPEEAIRRAATILAEQLEAFVDLRDVRQP
                 ************************************************************

RpoA_EDL399      EVKEEKPEFDPILLRPVDDLELTVRSANCLKAEAIHYIGDLVQRTEVELLKTPNLGKKSL
RpoA_W3110       EVKEEKPEFDPILLRPVDDLELTVRSANCLKAEAIHYIGDLVQRTEVELLKTPNLGKKSL
RpoA_042         EVKEEKPEFDPILLRPVDDLELTVRSANCLKAEAIHYIGDLVQRTEVELLKTPNLGKKSL
RpoA_MG1655      EVKEEKPEFDPILLRPVDDLELTVRSANCLKAEAIHYIGDLVQRTEVELLKTPNLGKKSL
RpoA_EMG2        EVKEEKPEFDPILLRPVDDLELTVRSANCLKAEAIHYIGDLVQRTEVELLKTPNLGKKSL
RpoA_WG1         EVKEEKPEFDPILLRPVDDLELTVRSANCLKAEAIHYIGDLVQRTEVELLKTPNLGKKSL
                 ************************************************************

                         311
RpoA_EDL399      TEIKDVLASRGLSLGMRLENWPPASIADE
RpoA_W3110       TEIKDVLASRGLSLGMRLENWPPASIADE
RpoA_042         TEIKDVLASRGLSLGMRLENWPPASIADE
RpoA_MG1655      TEIKDVLASRGLSLGMRLENWPPASIADE
RpoA_EMG2        TEIKDVLASRGLSLGMRLENWPPASIADE
RpoA_WG1         TEIKDVLASRRLSLGMRLENWPPASIADE
                 ********** ******************
```

### c) $\sigma^{38}$/ $\sigma^{S}$ (RpoS)

```
                                                 32
RpoS_W3110        MSQNTLKVHDLNEDAEFDENGVEVFDEKALVE----------------------------
RpoS_EMG2         MSQNTLKVHDLNEDAEFDENGVEVFDEKALVE----------------------------
RpoS_WG1          MSQNTLKVHDLNEDAEFDENGVEVFDEKALVE----------------------------
RpoS_EDL399       MSQNTLKVHDLNEDAEFDENGVEVFDEKALVEEEPSDNDLAEEELLSQGATQRVLDATQL
RpoS_MG1655       MSQNTLKVHDLNEDAEFDENGVEVFDEKALVEQEPSDNDLAEEELLSQGATQRVLDATQL
RpoS_BL21         MSQNTLKVHDLNEDAEFDENGVEVFDEKALVEEEPSDNDLAEEELLSQGATQRVLDATQL
RpoS_042          MSQNTLKVHDLNEDAEFDENGVEVFDEKALVEEEPSDNDLAEEELLSQGATQRVLDATQL
                  ********************************

RpoS_W3110        ------------------------------------------------------------
RpoS_EMG2         ------------------------------------------------------------
RpoS_WG1          ------------------------------------------------------------
RpoS_EDL399       YLGEIGYSPLLTAEEEVYFARRALRGDVASRRRMIESNLRLVVKIARRYGNRGLALLDLI
RpoS_MG1655       YLGEIGYSPLLTAEEEVYFARRALRGDVASRRRMIESNLRLVVKIARRYGNRGLALLDLI
RpoS_BL21         YLGEIGYSPLLTAEEEVYFARRALRGDVASRRRMIESNLRLVVKIARRYGNRGLALLDLI
RpoS_042          YLGEIGYSPLLTAEEEVYFARRALRGDVASRRRMIESNLRLVVKIARRYGNRGLALLDLI


RpoS_W3110        ------------------------------------------------------------
RpoS_EMG2         ------------------------------------------------------------
RpoS_WG1          ------------------------------------------------------------
RpoS_EDL399       EEGNLGLIRAVEKFDPERGFRFSTYATWWIRQTIERAIMNQTRTIRLPIHIVKELNVYLR
RpoS_MG1655       EEGNLGLIRAVEKFDPERGFRFSTYATWWIRQTIERAIMNQTRTIRLPIHIVKELNVYLR
RpoS_BL21         EEGNLGLIRAVEKFDPERGFRFSTYATWWIRQTIERAIMNQTRTIRLPIHIVKELNVYLR
RpoS_042          EEGNLGLIRAVEKFDPERGFRFSTYATWWIRQTIERAIMNQTRTIRLPIHIVKELNVYLR


RpoS_W3110        ------------------------------------------------------------
RpoS_EMG2         ------------------------------------------------------------
RpoS_WG1          ------------------------------------------------------------
RpoS_EDL399       TARELSHKLDHEPSAEEIAEQLDKPVDDVSRMLRLNERITSVDTPLGGDSEKALLDILAD
RpoS_MG1655       TARELSHKLDHEPSAEEIAEQLDKPVDDVSRMLRLNERITSVDTPLGGDSEKALLDILAD
RpoS_BL21         TARELSHKLDHEPSAEEIAEQLDKPVDDVSRMLRLNERITSVDTPLGGDSEKALLDILAD
RpoS_042          TARELSHKLDHEPSAEESAEQLDKPVDDVSRMLRLNERITSVDTPLGGDSEKALLDILAD


RpoS_W3110        ------------------------------------------------------------
RpoS_EMG2         ------------------------------------------------------------
RpoS_WG1          ------------------------------------------------------------
RpoS_EDL399       ------------------------------------------------------------
RpoS_MG1655       EKENGPEDTTQDDDMKQSIVKWLFELNAKQREVLARRFGLLGYEAATLEDVGREIGLTRE
RpoS_BL21         EKENGPEDTTQDDDMKQSIVKWLFELNAKQREVLARRFGLLGYEAATLEDVGREIGLTRE
RpoS_042          EKENGPEDTTQDDDMKQSIVKWLFELNAKQREVLARRFGLLGYEAATLEDVGREIGLTRE


RpoS_W3110        ------------------------------
RpoS_EMG2         ------------------------------
RpoS_WG1          ------------------------------
RpoS_EDL399       ------------------------------
RpoS_MG1655       RVRQIQVEGLRRLREILQTQGLNIEALFRE
RpoS_BL21         RVRQIQVEGLRRLREILQTQGLNIEALFRE
RpoS_042          RVRQIQVEGLRRLREILQTQGLNIEALFRE
```

### d) PrfB (RF2: Release factor 2)

```
PrfB_EMG2        MFEINPVNNRIQDLTERSDVLRGYLDYDAKKERLEEVNAELEQPDVWNEPERAQALGKER
PrfB_MG1655      MFEINPVNNRIQDLTERSDVLRGYLDYDAKKERLEEVNAELEQPDVWNEPERAQALGKER
PrfB_W3110       MFEINPVNNRIQDLTERSDVLRGYLDYDAKKERLEEVNAELEQPDVWNEPERAQALGKER
PrfB_WG1         MFEINPVNNRIQDLTERSDVLRGYLDYDAKKERLEEVNAELEQPDVWNEPERAQALGKER
PrfB_042         MFEINPVNNRIQDLTERSDVLRGYLDYDAKKERLEEVNAELEQPDVWNEPERAQALGKER
                 ************************************************************

PrfB_EMG2        SSLEAVVDTLDQMKQGLEDVSGLLELAVEADDEETFNEAVAELDALEEKLAQLEFRRMFS
PrfB_MG1655      SSLEAVVDTLDQMKQGLEDVSGLLELAVEADDEETFNEAVAELDALEEKLAQLEFRRMFS
PrfB_W3110       SSLEAVVDTLDQMKQGLEDVSGLLELAVEADDEETFNEAVAELDALEEKLAQLEFRRMFS
PrfB_WG1         SSLEAVVDTLDQMKQGLEDVSGLLELAVEADDEETFNEAVAELDALEEKLAQLEFRRMFS
PrfB_042         SSLEAVVDTLDQMKQGLEDVSGLLELAVEADDEETFNEAVAELDALEEKLAQLEFRRMFS
                 ************************************************************

PrfB_EMG2        GEYDSADCYLDIQAGSGGTEAQDWASMLERMYLRWAESRGFKTEIIEESEGEVAGIKSVT
PrfB_MG1655      GEYDSADCYLDIQAGSGGTEAQDWASMLERMYLRWAESRGFKTEIIEESEGEVAGIKSVT
PrfB_W3110       GEYDSADCYLDIQAGSGGTEAQDWASMLERMYLRWAESRGFKTEIIEESEGEVAGIKSVT
PrfB_WG1         GEYDSADCYLDIQAGSGGTEAQDWASMLERMYLRWAESRGFKTEIIEESEGEVAGIKSVT
PrfB_042         GEYDSADCYLDIQAGSGGTEAQDWASMLERMYLRWAESRGFKTEIIEESEGEVAGIKSVT
                 ************************************************************

PrfB_EMG2        IKISGDYAYGWLRTETGVHRLVRKSPFDSGGRRHTSFSSAFVYPEVDDDIDIEINPADLR
PrfB_MG1655      IKISGDYAYGWLRTETGVHRLVRKSPFDSGGRRHTSFSSAFVYPEVDDDIDIEINPADLR
PrfB_W3110       IKISGDYAYGWLRTETGVHRLVRKSPFDSGGRRHTSFSSAFVYPEVDDDIDIEINPADLR
PrfB_WG1         IKISGDYAYGWLRTETGVHRLVRKSPFDSGGRRHTSFSSAFVYPEVDDDIDIEINPADLR
PrfB_042         IKISGDYAYGWLRTETGVHRLVRKSPFDSGGRRHTSFSSAFVYPEVDDDIDIEINPADLR
                 ************************************************************
                    246
PrfB_EMG2        IDVYRTSGAGGQHVNRTESAVRITHIPTGIVTQCQNDRSQHKNKDQAMKQMKAKLYELEM
PrfB_MG1655      IDVYRTSGAGGQHVNRTESAVRITHIPTGIVTQCQNDRSQHKNKDQAMKQMKAKLYELEM
PrfB_W3110       IDVYRTSGAGGQHVNRTESAVRITHIPTGIVTQCQNDRSQHKNKDQAMKQMKAKLYELEM
PrfB_WG1         IDVYRASGAGGQHVNRTESAVRITHIPTGIVTQCQNDRSQHKNKDQAMKQMKAKLYELEM
PrfB_042         IDVYRASGAGGQHVNRTESAVRITHIPTGIVTQCQNDRSQHKNKDQAMKQMKAKLYELEM
                 *****:******************************************************

PrfB_EMG2        QKKNAEKQAMEDNKSDIGWGSQIRSYVLDDSRIKDLRTGVETRNTQAVLDGSLDQFIEAS
PrfB_MG1655      QKKNAEKQAMEDNKSDIGWGSQIRSYVLDDSRIKDLRTGVETRNTQAVLDGSLDQFIEAS
PrfB_W3110       QKKNAEKQAMEDNKSDIGWGSQIRSYVLDDSRIKDLRTGVETRNTQAVLDGSLDQFIEAS
PrfB_WG1         QKKNAEKQAMEDNKSDIGWGSQIRSYVLDDSRIKDLRTGVETRNTQAVLDGSLDQFIEAS
PrfB_042         QKKNAEKQAMEDNKSDIGWGSQIRSYVLDDSRIKDLRTGVETRNTQAVLDGSLDQFIEAS
                 ************************************************************

PrfB_EMG2        LKAGL
PrfB_MG1655      LKAGL
PrfB_W3110       LKAGL
PrfB_WG1         LKAGL
PrfB_042         LKAGL
                 *****
```

### e) RpsG

```
RpsG_WG1          MPRRRVIGQRKILPDPKFGSELLAKFVNILMVDGKKSTAESIVYSALETLAQRSGKSELE
RpsG_EMG2         MPRRRVIGQRKILPDPKFGSELLAKFVNILMVDGKKSTAESIVYSALETLAQRSGKSELE
RpsG_MG1655       MPRRRVIGQRKILPDPKFGSELLAKFVNILMVDGKKSTAESIVYSALETLAQRSGKSELE
RpsG_W3110        MPRRRVIGQRKILPDPKFGSELLAKFVNILMVDGKKSTAESIVYSALETLAQRSGKSELE
RpsG_EDL399       MPRRRVIGQRKILPDPKFGSELLAKFVNILMVDGKKSTAESIVYSALETLAQRSGKSELE
RpsG_042          MPRRRVIGQRKILPDPKFGSELLAKFVNILMVDGKKSTAESIVYSALETLAQRSGKSELE
                  ************************************************************

RpsG_WG1          AFEVALENVRPTVEVKSRRVGGSTYQVPVEVRPVRRNALAMRWIVEAARKRGDKSMALRL
RpsG_EMG2         AFEVALENVRPTVEVKSRRVGGSTYQVPVEVRPVRRNALAMRWIVEAARKRGDKSMALRL
RpsG_MG1655       AFEVALENVRPTVEVKSRRVGGSTYQVPVEVRPVRRNALAMRWIVEAARKRGDKSMALRL
RpsG_W3110        AFEVALENVRPTVEVKSRRVGGSTYQVPVEVRPVRRNALAMRWIVEAARKRGDKSMALRL
RpsG_EDL399       AFEVALENVRPTVEVKSRRVGGSTYQVPVEVRPVRRNALAMRWIVEAARKRGDKSMALRL
RpsG_042          AFEVALENVRPTVEVKSRRVGGSTYQVPVEVRPVRRNALAMRWIVEAARKRGDKSMALRL
                  ************************************************************

RpsG_WG1          ANELSDAAENKGTAVKKREDVHRMAEANKAFAHYRW-----------------------
RpsG_EMG2         ANELSDAAENKGTAVKKREDVHRMAEANKAFAHYRWLSLRSFSHQAGASSKQPALGYLN
RpsG_MG1655       ANELSDAAENKGTAVKKREDVHRMAEANKAFAHYRWLSLRSFSHQAGASSKQPALGYLN
RpsG_W3110        ANELSDAAENKGTAVKKREDVHRMAEANKAFAHYRWLSLRSFSHQAGASSKQPALGYLN
RpsG_EDL399       ANELSDAAENKGTAVKKREDVHRMAEANKAFAHYRW-----------------------
RpsG_042          ANELSDAAENKGTAVKKREDVHRMAEANKAFAHYRW-----------------------
                  ************************************
```

### f) Rph

```
Rph_MG1655       MRPAGRSNNQVRPVTLTRNYTKHAEGSVLVEFGDTKVLCTASIEEGVPRFLKGQGQGWIT
Rph_W3110        MRPAGRSNNQVRPVTLTRNYTKHAEGSVLVEFGDTKVLCTASIEEGVPRFLKGQGQGWIT
Rph_EMG2         MRPAGRSNNQVRPVTLTRNYTKHAEGSVLVEFGDTKVLCTASIEEGVPRFLKGQGQGWIT
Rph_EDL399       MRPAGRSNNQVRPVTLTRNYTKHAEGSVLVEFGDTKVLCTASIEEGVPRFLKGQGQGWIT
Rph_042          MRPAGRSNNQVRPVTLTRNYTKHAEGSVLVEFGDTKVLCTASIEEGVPRFLKGQGQGWIT
Rph_WG1          MRPAGRSNNQVRPVTLTRNYTKHAEGSVLVEFGDTKVLCTASIEEGVPRFLKGQGQGWIT
                 ************************************************************

Rph_MG1655       AEYGMLPRSTHTRNAREAAKGKQGGRTMEIQRLIARALRAAVDLKALGEFTITLDCDVLQ
Rph_W3110        AEYGMLPRSTHTRNAREAAKGKQGGRTMEIQRLIARALRAAVDLKALGEFTITLDCDVLQ
Rph_EMG2         AEYGMLPRSTHTRNAREAAKGKQGGRTMEIQRLIARALRAAVDLKALGEFTITLDCDVLQ
Rph_EDL399       AEYGMLPRSTHTRNAREAAKGKQGGRTMEIQRLIARALRAAVDLKALGEFTITLDCDVLQ
Rph_042          AEYGMLPRSTHTRNAREAAKGKQGGRTMEIQRLIARALRAAVDLKALGEFTITLDCDVLQ
Rph_WG1          AEYGMLPRSTHTRNAREAAKGKQGGRTMEIQRLIARALRAAVDLKALGEFTITLDCDVLQ
                 ************************************************************

Rph_MG1655       ADGGTRTASITGACVALVDALQKLVENGKLKTNPMKGMVAAVSVGIVNGEAVCDLEYVED
Rph_W3110        ADGGTRTASITGACVALVDALQKLVENGKLKTNPMKGMVAAVSVGIVNGEAVCDLEYVED
Rph_EMG2         ADGGTRTASITGACVALVDALQKLVENGKLKTNPMKGMVAAVSVGIVNGEAVCDLEYVED
Rph_EDL399       ADGGTRTASITGACVALADALQKLVENGKLKTNPMKGMVAAVSVGIVNGEAVCDLEYVED
Rph_042          ADGGTRTASITGACVALADALQKLVENGKLKTNPMKGMVAAVSVGIVNGEAVCDLEYVED
Rph_WG1          ADGGTRTASITGACVALVDALQKLVENGKLKTNPMKGMVAAVSVGIVNGEAVCDLEYVED
                 *****************.******************************************

Rph_MG1655       SAAETDMNVVMTEDGRIIEVQGTAEGEPFTHEELLILLALARGESNPL----------
Rph_W3110        SAAETDMNVVMTEDGRIIEVQGTAEGEPFTHEELLILLALARGESNPL----------
Rph_EMG2         SAAETDMNVVMTEDGRIIEVQGTAEGEPFTHEELLILLALARGESNPL----------
Rph_EDL399       SAAETDMNVVMTEDGRIIEVQGTAEGEPFTHEELLTLLALARGGIESIVATQKAALAN
Rph_042          SAAETDMNVVMTEDGRIIEVQGTAEGEPFTHEELLTLLALARGGIESIVATQKAALAN
Rph_WG1          SAAETDMNVVMTEDGRIIEVQGTAEGEPFTHEELLILLALARGGIESIVATQKAALAN
                 *********************************** *******  :.:
```

### g) IlvG

```
IlvG_WG1         TGKTGVCIATSGPGATNLITGLADALLDSIPVVAITGQVSAPFIGTDAFQEVDVLGLSLA
IlvG_EDL399      TGKTGVCIATSGPGATNLITGLADALLDSIPVVAITGQVSAPFIGTDAFQEVDVLGLSLA
IlvG_042         TGKTGVCIATSGPGATNLITGLADALLDSIPVVAITGQVSAPFIGTDAFQEVDVLGLSLA
IlvG_EMG2        TGKTGVCIATSGPGATNLITGLADALLDSIPVVAITGQVSAPFIGTDAFQEVDVLGLSLA
                 ************************************************************

IlvG_WG1         CTKHSFLVQSLEELPRIMAEAFDVACSGRPGPVLVDIPKDIQLASGDLEPWFTTVENEVT
IlvG_EDL399      CTKHSFLVQSLEELPRIMAEAFDVASSGRPGPVLVDIPKDIQLASGDLEPWFTTVENEVT
IlvG_042         CTKHSFLVQSLEELPRIMAEAFDVAGSGRPGPVLVDIPKDIQLASGDLEPWFTTVENEVN
IlvG_EMG2        CTKHSFLVQSLEELPRIMAEAFDVACSGRPGPVLVDIPKDIQLASGDLEPWFTTVENEVT
                 ************************* *********************************.

IlvG_WG1         FPHAEVEQARQMLAKAQKPMLYVGGGVGMAQAVPALREFLAATKMPATCTLKGLGAVEAD
IlvG_EDL399      FPHAEVEQARQMLAKAQKPMLYVGGGVGMAQAVPALREFLATTKMPATCTLKGLGAVEAD
IlvG_042         FPHAEVEQARQMLAKAQKPMLYVGGGVGMAQAVPALREFLAATKMPVTCTLKGLGAVEAD
IlvG_EMG2        FPHAEVEQARQMLAKAQKPMLYVGGGVGMAQAVPALREFLAATKMPATCTLKGLGAVEAD
                 *****************************************:****.*************

IlvG_WG1         YPYYLGMLGMHGTKAANFAVQECDLLIAVGARFDDRVTGKLNTFAPHASVIHMDIDPAEM
IlvG_EDL399      YPYYLGMLGMHGTKAANFAVQECDLLIAVGARFDDRVTGKLNTFAPHASVIHMDIDPAEM
IlvG_042         YPYYLGMLGMHGTKAANFAVQECDLLIAVGARFDDRVTGKLNTFAPLASVIHMDIDPAEM
IlvG_EMG2        YPYYLGMLGMHGTKAANFAVQECDLLIAVGARFDDRVTGKLNTFAPHASVIHMDIDPAEM
                 ********************************************** *************

IlvG_WG1         NKLRQAHVALQGDLNALLPALQQPLNINDWQQHCAQLRDEHSWRYDHPGDAIYAPLLLKQ
IlvG_EDL399      NKLRQAHVALQGDLNALLPALQQPLNINDWQQYCAQLRDEHAWRYDHPGDAIYAPLLLKQ
IlvG_042         NKLRQAHVALQGDLNALLPALQQPLNINDWQQYCAQLRDEHTWRYDHPGDAIYAPLLLKQ
IlvG_EMG2        NKLRQAHVALQGDLNALLPALQQPLNQ---------------------------------
                 **************************

IlvG_WG1         LSDRKPADCVVTTDVGQHQMWAAQHIAHTRPENFITSSGLGTMGFGLPAAVGAQVARPND
IlvG_EDL399      LSDRKPADCVVTTDVGQHQMWAAQHIAHTRPENFITSSGLGTMGFGLPAAVGAQVARPND
IlvG_042         LSDRKPADCVVTTDVGQHQMWAAQHIAHTRPENFITSSGLGTMGFGLPAAVGAQVARPND
IlvG_EMG2        ------------------------------------------------------------

IlvG_WG1         TVVCISGDGSFMMNVQELGTVKRKQLPLKIVLLDNQRLGMVRQWQQLFFQERYSETTLTD
IlvG_EDL399      TVVCISGDGSFMMNVQELGTVKRKQLPLKIVLLDNQRLGMVRQWQQLFFQERYSETTLTD
IlvG_042         TVVCISGDGSFMMNVQELGTVKRKQLPLKIVLLDNQRLGMVRQWQQLFFQERYSETTLTD
IlvG_EMG2        ------------------------------------------------------------

IlvG_WG1         NPDFLMLASAFGIHGQHITRKDQVEAALDTMLNSDGPYLLHVSIDELENVWPLVPPGASN
IlvG_EDL399      NPDFLKLASAFGIPGQHITRKDQVEAALDTMLNSDGPYLLHVSIDELENVWPLVPPGASN
IlvG_042         NPDFLMLASAFGIPGQHITRKDQVEAALDTMLNSDGPYLLHVSIDELENVWPLVPPGASN
IlvG_EMG2        ------------------------------------------------------------

IlvG_WG1         SEMLEKLS
IlvG_EDL399      SEMLEKLS
IlvG_042         SEMLEKLS
IlvG_EMG2        --------
```

### h) MdtF

```
MdtF_WG1         MANYFIDRPVFAWVLAIIMMLAGGLAIMNLPVAQYPQIAPPTITVSATYPGADAQTVEDS
MdtF_EMG2        MANYFIDRPVFAWVLAIIMMLAGGLAIMNLPVAQYPQIAPPTITVSATYPGADAQTVEDS
MdtF_MG1655      MANYFIDRPVFAWVLAIIMMLAGGLAIMNLPVAQYPQIAPPTITVSATYPGADAQTVEDS
MdtF_W3110       MANYFIDRPVFAWVLAIIMMLAGGLAIMNLPVAQYPQIAPPTITVSATYPGADAQTVEDS
MdtF_EDL399      MANYFIDRPVFAWVLAIIMMLAGGLAIMNLPVAQYPQIAPPTITVSATYPGADAQTVEDS
MdtF_042         MANYFIDRPVFAWVLAIIMMLAGGLAIMNLPVAQYPQIAPPTITISATYPGADAQTVEDS
                 ********************************************:***************
```

```
MdtF_WG1        VTQVIEQNMNGLDGLMYMSSTSDAAGNASITLTFETGTSPDIAQVQVQNKLQLAMPSLPE
MdtF_EMG2       VTQVIEQNMNGLDGLMYMSSTSDAAGNASITLTFETGTSPDIAQVQVQNKLQLAMPSLPE
MdtF_MG1655     VTQVIEQNMNGLDGLMYMSSTSDAAGNASITLTFETGTSPDIAQVQVQNKLQLAMPSLPE
MdtF_W3110      VTQVIEQNMNGLDGLMYMSSTSDAAGNASITLTFETGTSPDIAQVQVQNKLQLAMPSLPE
MdtF_EDL399     VTQVIEQNMNGLDGLMYMSSTSDAAGNASITLTFETGTSPDIAQVQVQNKLQLAMPSLPE
MdtF_042        VTQVIEQNMNGLDGLMYMSSTSDAAGNASITLTFETGTSPDIAQVQVQNKLQLAMPSLPE
                ************************************************************

MdtF_WG1        AVQQQGISVDKSSSNILMVAAFISDNGSLNQYDIADYVASNIKDPLSRTAGVGSVQLFGS
MdtF_EMG2       AVQQQGISVDKSSSNILMVAAFISDNGSLNQYDIADYVASNIKDPLSRTAGVGSVQLFGS
MdtF_MG1655     AVQQQGISVDKSSSNILMVAAFISDNGSLNQYDIADYVASNIKDPLSRTAGVGSVQLFGS
MdtF_W3110      AVQQQGISVDKSSSNILMVAAFISDNGSLNQYDIADYVASNIKDPLSRTAGVGSVQLFGS
MdtF_EDL399     AVQQQGISVDKSSSNILMVAAFISDNGSLNQYDIADYVASNIKDPLSRTAGVGSVQLFGS
MdtF_042        AVQQQGISVDKSSSNILMVAAFISDNGSLNQYDIADYVASNIKDPLSRTAGVGSVQLFGS
                ************************************************************

MdtF_WG1        EYAMRIWLDPQKLNKYNLVPSDVISQIKVQNNQISGGQLGGMPQAADQQLNASIIVQTRL
MdtF_EMG2       EYAMRIWLDPQKLNKYNLVPSDVISQIKVQNNQISGGQLGGMPQAADQQLNASIIVQTRL
MdtF_MG1655     EYAMRIWLDPQKLNKYNLVPSDVISQIKVQNNQISGGQLGGMPQAADQQLNASIIVQTRL
MdtF_W3110      EYAMRIWLDPQKLNKYNLVPSDVISQIKVQNNQISGGQLGGMPQAADQQLNASIIVQTRL
MdtF_EDL399     EYAMRIWLDPQKLNKYNLVPSDVISQIKVQNNQISGGQLGGMPQAADQQLNASIIVQTRL
MdtF_042        EYAMRIWLDPQKLNKYNLVPSDVISQIKVQNNQISGGQLGGMPQAADQQLNASIIVQTRL
                ************************************************************

MdtF_WG1        QTPEEFGKILLKVQQDGSQVLLRDVARVELGAEDYSTVARYNGKPAAGIAIKLAAGANAL
MdtF_EMG2       QTPEEFGKILLKVQQDGSQVLLRDVARVELGAEDYSTVARYNGKPAAGIAIKLAAGANAL
MdtF_MG1655     QTPEEFGKILLKVQQDGSQVLLRDVARVELGAEDYSTVARYNGKPAAGIAIKLAAGANAL
MdtF_W3110      QTPEEFGKILLKVQQDGSQVLLRDVARVELGAEDYSTVARYNGKPAAGIAIKLAAGANAL
MdtF_EDL399     QTPEEFGKILLKVQQDGSQVLLRDVARVELGAEDYSTVARYNGKPAAGIAIKLATGANAL
MdtF_042        QTPEEFGKILLKVQQDGSQVLLRDVARVELGAEDYSTVARYNGKPAAGIAIKLATGANAL
                ******************************************************:*****

MdtF_WG1        DTSRAVKEELNRLSAYFPASLKTVYPYDTTPFIEISIQEVFKTLVEAIILVFLVMYLFLQ
MdtF_EMG2       DTSRAVKEELNRLSAYFPASLKTVYPYDTTPFIEISIQEVFKTLVEAIILVFLVMYLFLQ
MdtF_MG1655     DTSRAVKEELNRLSAYFPASLKTVYPYDTTPFIEISIQEVFKTLVEAIILVFLVMYLFLQ
MdtF_W3110      DTSRAVKEELNRLSAYFPASLKTVYPYDTTPFIEISIQEVFKTLVEAIILVFLVMYLFLQ
MdtF_EDL399     DTSRAVKEELNRLSAYFPASLKTVYPYDTTPFIEISIQEVFKTLVEAIILVFLVMYLFLQ
MdtF_042        DTSRAVKEELNRLSAYFPASLKTVYPYDTTPFIEISIQEVFKTLVEAIILVFLVMYLFLQ
                ************************************************************

MdtF_WG1        NFRATIIPTIAVPVVILGTFAILSAVGFTINTLTMFGMVLAIGLLVDDAIVVVENVERVI
MdtF_EMG2       NFRATIIPTIAVPVVILGTFAILSAVGFTINTLTMFGMVLAIGLLVDDAIVVVENVERVI
MdtF_MG1655     NFRATIIPTIAVPVVILGTFAILSAVGFTINTLTMFGMVLAIGLLVDDAIVVVENVERVI
MdtF_W3110      NFRATIIPTIAVPVVILGTFAILSAVGFTINTLTMFGMVLAIGLLVDDAIVVVENVERVI
MdtF_EDL399     NFRATIIPTIAVPVVILGTFAILSAVGFTINTLTMFGMVLAIGLLVDDAIVVVENVERVI
MdtF_042        NFRATIIPTIAVPVVILGTFAILSAVGFTINTLTMFGMVLAIGLLVDDAIVVVENVERVI
                ************************************************************

MdtF_WG1        AEDKLPPKEATHKSMGQIQRALVGIAVVLSAVFMPMAFMSGATGEIYRQFSITLISSMLL
MdtF_EMG2       AEDKLPPKEATHKSMGQIQRALVGIAVVLSAVFMPMAFMSGATGEIYRQFSITLISSMLL
MdtF_MG1655     AEDKLPPKEATHKSMGQIQRALVGIAVVLSAVFMPMAFMSGATGEIYRQFSITLISSMLL
MdtF_W3110      AEDKLPPKEATHKSMGQIQRALVGIAVVLSAVFMPMAFMSGATGEIYRQFSITLISSMLL
MdtF_EDL399     AEDKLPPKEATHKSMGQIQRALVGIAVVLSAVFMPMAFMSGATGEIYRQFSITLISSMLL
MdtF_042        AEDKLPPKEATHKSMGQIQRALVGIAVVLSAVFMPMAFMSGATGEIYRQFSITLISSMLL
                ************************************************************

MdtF_WG1        SVFVAMSLTPALCATILKAAPEGGHKPNALFARFNTLFEKSTQHYTDSTRSLLRCTGRYM
MdtF_EMG2       SVFVAMSLTPALCATILKAAPEGGHKPNALFARFNTLFEKSTQHYTDSTRSLLRCTGRYM
MdtF_MG1655     SVFVAMSLTPALCATILKAAPEGGHKPNALFARFNTLFEKSTQHYTDSTRSLLRCTGRYM
MdtF_W3110      SVFVAMSLTPALCATILKAAPEGGHKPNALFARFNTLFEKSTQHYTDSTRSLLRCTGRYM
MdtF_EDL399     SVFVAMSLTPALCATILKAAPEGGHKPNALFARFNTLFEKSTQHYTDSTRSLLRCTGRYM
MdtF_042        SVFVAMSLTPALCATILKAAPEGGHKPNALFARFNTLFEKSTQHYTNSTRSLLRCTGRYM
                **********************************************:*************
```

```
MdtF_WG1        VVYLLICAGMAVLFLRTPTSFLPEEDQGVFMTTAQLPSGATMVNTTKVLQQVTDYYLTKE
MdtF_EMG2       VVYLLICAGMAVLFLRTPTSFLPEEDQGVFMTTAQLPSGATMVNTTKVLQQVTDYYLTKE
MdtF_MG1655     VVYLLICAGMAVLFLRTPTSFLPEEDQGVFMTTAQLPSGATMVNTTKVLQQVTDYYLTKE
MdtF_W3110      VVYLLICAGMAVLFLRTPTSFLPEEDQGVFMTTAQLPSGATMVNTTKVLQQVTDYYLTKE
MdtF_EDL399     VIYLLICAGMAVLFLRTPTSFLPEEDQGVFMTTAQLPSGATMVNTTKVLQQVTDYYLTKE
MdtF_042        VVYLLICAGMAVLFLRTPTSFLPEEDQGVFMTTAQLPSGATMVNTTKVLQQVTDYYLTKE
                *:**********************************************************

MdtF_WG1        KDNVQSVFTVGGFGFSGQGQNNGLAFISLKPWSERVGEENSVTAIIQRAMIALSSINKAV
MdtF_EMG2       KDNVQSVFTVGGFGFSGQGQNNGLAFISLKPWSERVGEENSVTAIIQRAMIALSSINKAV
MdtF_MG1655     KDNVQSVFTVGGFGFSGQGQNNGLAFISLKPWSERVGEENSVTAIIQRAMIALSSINKAV
MdtF_W3110      KDNVQSVFTVGGFGFSGQGQNNGLAFISLKPWSERVGEENSVTAIIQRAMIALSSINKAV
MdtF_EDL399     KDNVQSVFTVGGFGFSGQGQNNGLAFISLKPWSERVGEENSVTAIIQRAMIALSSINKAV
MdtF_042        KDNVQSVFTVGGFGFSGQGQNNGLAFISLKPWSERVGEENSVTAIIQRAMIALSSINKAV
                ************************************************************

MdtF_WG1        VFPFNLPAVAELGTASGFDMELLDNGNLGHEKLTQARNELLSLAAQSPNQVTGVRPNGLE
MdtF_EMG2       VFPFNLPAVAELGTASGFDMELLDNGNLGHEKLTQARNELLSLAAQSPNQVTGVRPNGLE
MdtF_MG1655     VFPFNLPAVAELGTASGFDMELLDNGNLGHEKLTQARNELLSLAAQSPNQVTGVRPNGLE
MdtF_W3110      VFPFNLPAVAELGTASGFDMELLDNGNLGHEKLTQARNELLSLAAQSPNQVTGVRPNGLE
MdtF_EDL399     VFPFNLPAVAELGTASGFDMELLDNGNLGHEKLTQARNELLSLAAQSPNQVTGVRPNGLE
MdtF_042        VFPFNLPAVAELGTASGFDMELLDNGNLGHEKLTQARNELLSLAAQSPNQVTGVRPNGLE
                ************************************************************

MdtF_WG1        DTPMFKVNVNAAKAEAMGVALSDINQTISTAFGSSYVNDFLN------------------
MdtF_EMG2       DTPMFKVNVNAAKAEAMGVALSDINQTISTAFGSSYVNDFLNQGRVKKVYVQAGTPFRML
MdtF_MG1655     DTPMFKVNVNAAKAEAMGVALSDINQTISTAFGSSYVNDFLNQGRVKKVYVQAGTPFRML
MdtF_W3110      DTPMFKVNVNAAKAEAMGVALSDINQTISTAFGSSYVNDFLNQGRVKKVYVQAGTPFRML
MdtF_EDL399     DTPMFKVNVNAAKAEAMGVALSDINQTISTAFGSSYVNDFLNQGRVKKVYVQAGTPFRML
MdtF_042        DTPMFKVNVNAAKAEAMGVALSDINQTISTAFGSSYVNDFLNQGRVKKVYVQAGTPFRML
                ******************************************
MdtF_WG1        ------------------------------------------------------------
MdtF_EMG2       PDNINQWYVRNASGTMAPLSAYSSTEWTYGSPRLERYNGIPSMEILGEAAAGKSTGDAMK
MdtF_MG1655     PDNINQWYVRNASGTMAPLSAYSSTEWTYGSPRLERYNGIPSMEILGEAAAGKSTGDAMK
MdtF_W3110      PDNINQWYVRNASGTMAPLSAYSSTEWTYGSPRLERYNGIPSMEILGEAAAGKSTGDAMK
MdtF_EDL399     PDNINQWYVRNASGTMAPLSAYSSTEWTYGSPRLERYNGIPSMEILGEAAAGKSTGDAMK
MdtF_042        PDNINQWYVRNASGTMAPLSAYSSTEWTYGSPRLERYNGIPSMEILGEAAAGKSTGDAMK

MdtF_WG1        ------------------------------------------------------------
MdtF_EMG2       FMADLVAKLPAGVGYSWTGLSYQEALSSNQAPALYAISLVVVFLALAALYESWSIPFSVM
MdtF_MG1655     FMADLVAKLPAGVGYSWTGLSYQEALSSNQAPALYAISLVVVFLALAALYESWSIPFSVM
MdtF_W3110      FMADLVAKLPAGVGYSWTGLSYQEALSSNQAPALYAISLVVVFLALAALYESWSIPFSVM
MdtF_EDL399     FMADLVAKLPAGVGYSWTGLSYQEALSSNQAPALYAISLVVVFLALAALYESWSIPFSVM
MdtF_042        FMADLVAKLPAGVGYSWTGLSYQEALSSNQAPALYAISLVVVFLALAALYESWSIPFSVM

MdtF_WG1        ------------------------------------------------------------
MdtF_EMG2       LVVPLGVVGALLATDLRGLSNDVYFQVGLLTTIGLSAKNAILIVEFAVEMMQKEGKTPIE
MdtF_MG1655     LVVPLGVVGALLATDLRGLSNDVYFQVGLLTTIGLSAKNAILIVEFAVEMMQKEGKTPIE
MdtF_W3110      LVVPLGVVGALLATDLRGLSNDVYFQVGLLTTIGLSAKNAILIVEFAVEMMQKEGKTPIE
MdtF_EDL399     LVVPLGVVGALLATDLRGLSNDVYFQVGLLTTIGLSAKNAILIVEFAVEMMQKEGKTPIE
MdtF_042        LVVPLGVVGALLATDLRGLSNDVYFQVGLLTTIGLSAKNAILIVEFAVEMMQKEGKTPIE


MdtF_WG1        ------------------------------------------------------------
MdtF_EMG2       AIIEAARMRLRPILMTSLAFILGVLPLVISHGAGSGAQNAVGTGVMGGMFAATVLAIYFV
MdtF_MG1655     AIIEAARMRLRPILMTSLAFILGVLPLVISHGAGSGAQNAVGTGVMGGMFAATVLAIYFV
MdtF_W3110      AIIEAARMRLRPILMTSLAFILGVLPLVISHGAGSGAQNAVGTGVMGGMFAATVLAIYFV
MdtF_EDL399     AIIEAARMRLRPILMTSLAFILGVLPLVISHGAGSGAQNAVGTGVMGGMFAATVLAIYFV
MdtF_042        AIIEAARMRLRPILMTSLAFILGLLPLVISHGAGSGAQNAVGTGVMGGMFAATVLAIYFV
```

```
MdtF_WG1          -----------------
MdtF_EMG2         PVFFVVVEHLFARFKKA
MdtF_MG1655       PVFFVVVEHLFARFKKA
MdtF_W3110        PVFFVVVEHLFARFKKA
MdtF_EDL399       PVFFVVVEHLFARFKKA
MdtF_042          PVFFVVVEHLFARFKKA
```

### i) Nfi

```
Nfi_EMG2          MDLASLRAQQIELASSVIREDRLDKDPPDLIAGADVGFEQGGEVTRAAMVLLKYPSLELV
Nfi_042           MDLASLRAQQIELASSVIREDRLDKDPPDLIAGADVGFEQGGEVTRAAMVLLNYPSLELV
Nfi_MG1655        MDLASLRAQQIELASSVIREDRLDKDPPDLIAGADVGFEQGGEVTRAAMVLLKYPSLELV
Nfi_EDL399        MDLASLRAQQIELASSVIREDRLDKDPPDLIAGADVGFEQGGEVTRAAMVLLKYPSLELV
Nfi_W3110         MDLASLRAQQIELASSVIREDRLDKDPPDLIAGADVGFEQGGEVTRAAMVLLKYPSLELV
Nfi_WG1           MDLASLRAQQIELASSVIREDRLDKDPPDLIAGADVGFEQGGEVTRAAMVLLKYPSLELV
                  ****************************************************:*******

Nfi_EMG2          EYKVARIATTMPYIPGFLSFREYPALLAAWEMLSQKPDLVFVDGHGISHPRRLGVASHFG
Nfi_042           EYKVARIATTMPYIPGFLSFREYPALLAAWEMLSQKPDLVFVDGHGISHPRRLGVASHFG
Nfi_MG1655        EYKVARIATTMPYIPGFLSFREYPALLAAWEMLSQKPDLVFVDGHGISHPRRLGVASHFG
Nfi_EDL399        EYKVARIATTMPYIPGFLSFREYPALLAAWEMLSQKPDLVFVDGHGISHPRRLGVASHFG
Nfi_W3110         EYKVARIATTMPYIPGFLSFREYPALLAAWEMLSQKPDLVFVDGHGISHPRRLGVASHFG
Nfi_WG1           EYKVARIATTMPYIPGFLSFREYPALLAAWEMLSQKPDLVFVDGHGISHPRRLGVASHFG
                  ************************************************************

Nfi_EMG2          LLVDVPTIGVAKKRLCGKFEPLSSEPGALAPLMDKGEQLAWVWRSKARCNPLFIATGHRV
Nfi_042           LLVDVPTIGVAKKRLCGKFEPLSSEPGALAPLMDKGEQLAWVWRSKARCNPLFIATGHRV
Nfi_MG1655        LLVDVPTIGVAKKRLCGKFEPLSSEPGALAPLMDKGEQLAWVWRSKARCNPLFIATGHRV
Nfi_EDL399        LLVDVPTIGVAKKRLCGKFEPLSSEPGALAPLMDKGEQLAWVWRSKARCNPLFIATGHRV
Nfi_W3110         LLVDVPTIGVAKKRLCGKFEPLSSEPGALAPLMDKGEQLAWVWRSKARCNPLFIATGHRV
Nfi_WG1           LLVDVPTIGVAKKRLCGKFEPLSSEPGALAPLMDKGEQLAWVWRAAKRAVTRCLSLPAIG
                  ********************************************:  *. .   ::

Nfi_EMG2          SVDSALAWVQRCMKGYRLPEPTRWADAVASERPAFVRYTANQP
Nfi_042           SVDSALAWVQRCMKGYRLPEPTRWADAVASERPAFVRYTANQP
Nfi_MG1655        SVDSALAWVQRCMKGYRLPEPTRWADAVASERPAFVRYTANQP
Nfi_EDL399        SVDSALAWVQRCMKGYRLPEPTRWADAVASERPAFVRYTANQP
Nfi_W3110         SVDSALAWVQRCMKGYRLPEPTRWADAVASERPAFVRYTANQP
Nfi_WG1           SAWTARWRGYNAA------------------------------
                  *. :*     ..
```

**Supplementary Fig. S13.**

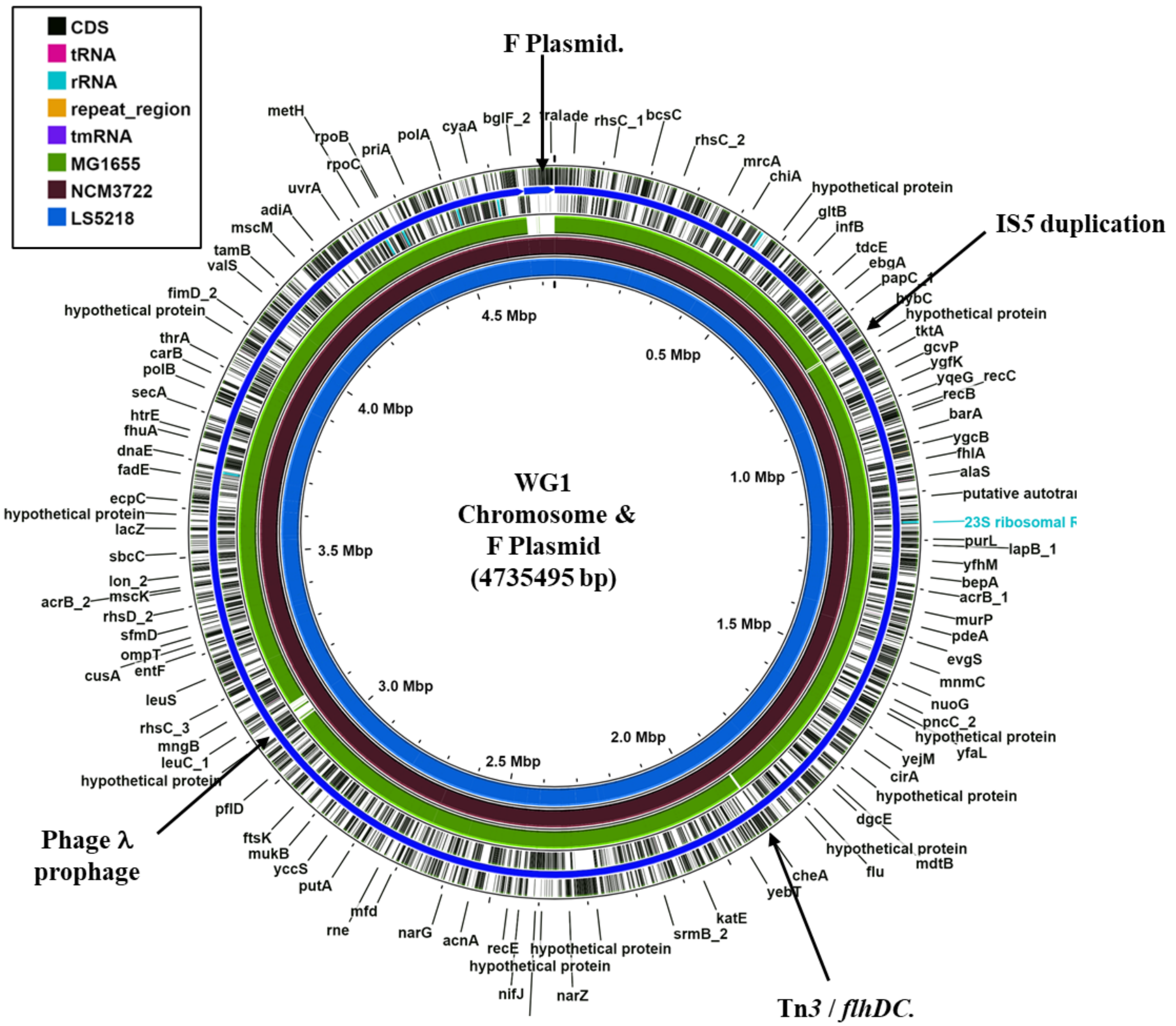

**Supplementary Fig. S14.**

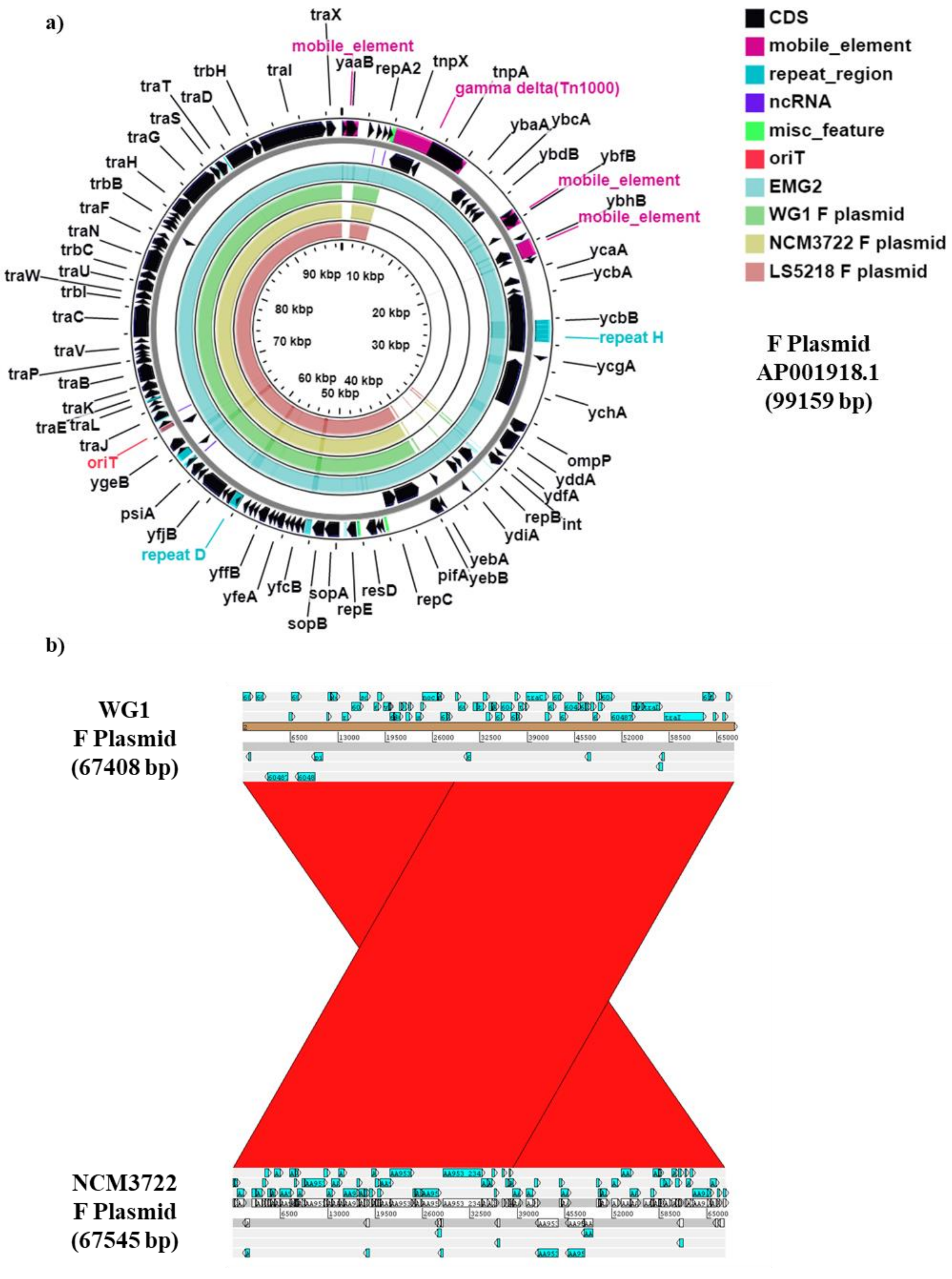
